# Heterogeneity in human hippocampal CaMKII transcripts reveals allosteric hub-dependent regulation

**DOI:** 10.1101/721589

**Authors:** Roman Sloutsky, Noelle Dziedzic, Matthew J. Dunn, Rachel M. Bates, Ana P. Torres-Ocampo, Sivakumar Boopathy, Brendan Page, John G. Weeks, Luke H. Chao, Margaret M. Stratton

## Abstract

Ca^2+^-calmodulin dependent protein kinase II (CaMKII) plays a central role in Ca^2+^ signaling throughout the body. Specifically in the hippocampus, CaMKII is required for learning and memory. CaMKII is encoded by four highly conserved genes in vertebrates: α, β, γ, and δ. All CaMKIIs are comprised of a kinase domain, regulatory segment, variable linker region, and hub domain responsible for oligomerization. The four genes differ primarily in linker length and composition due to extensive alternative splicing. Here, we unambiguously report the heterogeneity of CaMKII transcripts in 3 complex samples of human hippocampus using Illumina sequencing. Our results show that hippocampal cells contain a diverse collection of 70 CaMKII transcripts from all four CaMKII genes. We characterized the Ca^2+^/CaM sensitivity of hippocampal CaMKII variants spanning a broad range of linker lengths and compositions. We demonstrate that the effect of the variable linker on Ca^2+^/CaM sensitivity is conditional on kinase and hub domains. Moreover, we reveal a novel role for the hub domain as an allosteric regulator of kinase activity, which may provide a new pharmacological target for modulating CaMKII activity. Using small angle X-ray scattering and single-particle electron cryo-microscopy, we present evidence for extensive interaction between the kinase and the hub domain, even in the presence of a 30-residue linker. Taken together, we propose that Ca^2+^/CaM sensitivity in CaMKII is gene-dependent and includes significant contributions from the hub. Our sequencing approach combined with biochemistry provides new insights into understanding the complex pool of endogenous CaMKII.

**One Sentence Summary:** CaMKII is a well-conserved protein that is essential for learning and memory. When CaMKII is mutated in a mouse, this mouse has difficulty learning and remembering how to get through a maze. The hippocampus is the part of the brain required for memory. Here, we used a specific experiment to determine every type of CaMKII that is in a human hippocampus. We found 70 different types and then asked how these differences affect CaMKII function. These data provide evidence that an assembly domain of CaMKII plays an unexpected role regulating its activity. This new finding helps us better understand endogenous CaMKII in the brain and provides a new mechanism for modulating CaMKII activity.

## Introduction

Ca^2+^/calmodulin dependent protein kinase II (CaMKII) is a crucial oligomeric Ser/Thr kinase in long-term memory formation (*1*) as well as egg activation in fertilization (*2, 3*), and cardiac pacemaking (*4*). CaMKII is expressed throughout the human body, which facilitates its role in these disparate functions. There are four human genes that encode CaMKII: CaMKIIα and β (predominantly in brain), CaMKIIδ (predominant in heart) and CaMKIIγ (multiple organ systems, including egg and sperm) (*5–7*). In all four genes, each CaMKII subunit is comprised of a kinase domain, regulatory segment, variable linker region, and a hub domain (Fig. 1A). The kinase and hub domains are highly conserved across the four genes, with minimum 90% and 75% pairwise identity, respectively. The linker that connects the kinase and hub domains is highly variable in length and composition due to alternative splicing. Herein, we focus on CaMKII in the hippocampus, the primary memory center in the brain (*8*). CaMKII is the most abundant enzyme in neuronal dendrites and has been implicated functionally in memory (*1*).

**Figure 1.**
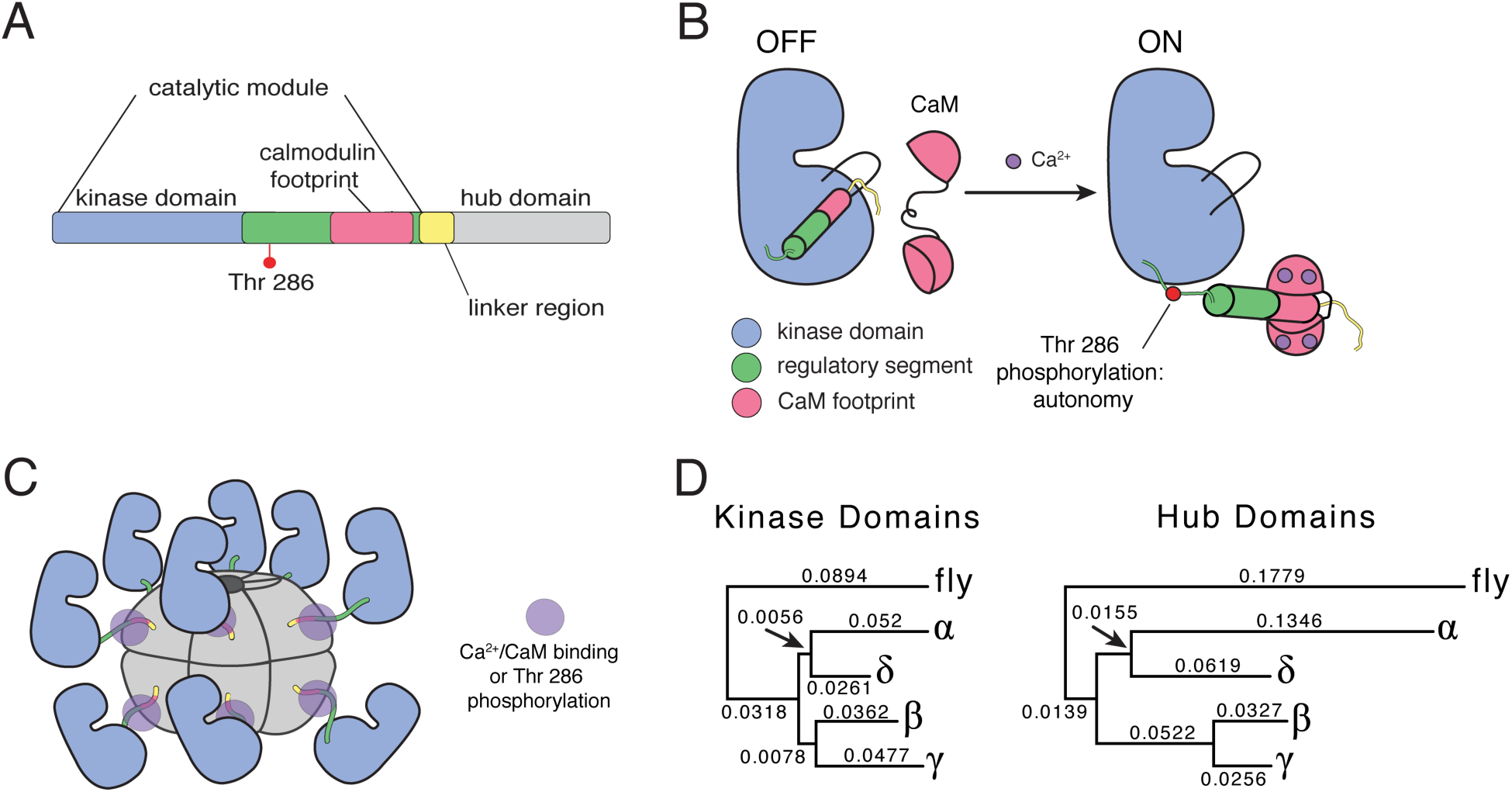
CaMKII sequence divergence and structural organization. (**A**) Each subunit of CaMKII consists of a kinase domain, regulatory segment, variable linker region, and hub domain. The regulatory segment houses the calmodulin binding region as well as a regulatory autophosphorylation site, Thr^286^. (**B**) The regulatory segment acts as an autoinhibitory domain by blocking the substrate-binding pocket. In the presence of Ca^2+^, Ca^2+^-CaM binds the regulatory segment, exposing the substrate-binding pocket and turning the enzyme on. (**C**) The hub domain oligomerizes CaMKII into two stacked hexameric rings. Shown is a cartoon of an active structure with Ca^2+^-CaM bound to the regulatory segment. (**D**) Neighbor Joining reconstructions of kinase and hub domain divergence for the four human CaMKII genes, rooted with *D. melanogaster* CaMKII outgroups.

Transgenic mice deficient in neuronal CaMKII have limited long-term memory and display specific learning impairments (*9, 10*). More specifically, mice with mutations at critical phosphorylation sites in CaMKII display defects in higher order memory and adaptive learning (*11*).

The structure and function of both the kinase and hub domains of CaMKII have been well studied. Kinase activity is regulated by the autoinhibitory regulatory segment, which blocks the substrate-binding site in the absence of Ca^2+^ (*12*). When Ca^2+^ levels rise, Ca^2+^ bound calmodulin (Ca^2+^/CaM) competitively binds to the regulatory segment and relieves inhibition by exposing the substrate-binding site (Fig. 1B) (*13*). The regulatory segment also contains three sites of autophosphorylation: Thr^286^, Thr^305^, Thr^306^ (following CaMKIIα numbering) (*14*). Notably, phosphorylation at Thr^286^ results in autonomous activation, or, sustained activity in the absence of Ca^2+^ (*15*). The hub domain oligomerizes the holoenzyme into both dodecameric and tetradecameric assemblies (Fig. 1C) (*16–18*).

The variable linker region has been difficult to study for several reasons. The sequence of the linker is predicted to be disordered. It is likely for this reason that high-resolution structures of the CaMKII holoenzyme including the linker region have not been solved. Indeed, the only atomic resolution structure of a CaMKII holoenzyme is one in which the variable linker is completely deleted (*19*). In this structure, the regulatory segment makes extensive contacts with the hub domain, thereby completely sequestering the CaM binding segment from Ca^2+^/CaM in this compact conformation. This structure, combined with biochemical data discussed below, led to the hypothesis that linker length tunes the autoinhibition within the holoenzyme by either stabilizing a compact conformation (short linker forms) or populating an extended conformation where the CaM binding segment is more accessible (longer linker forms).

The variable linker also varies in composition between the four CaMKII genes. Further complexity is generated by alternative splicing of up to 9 exons (exons 13-21) in each linker, creating a total of more than 50 variants possibly expressed from four genes (Fig. 2A). The combination of high identity within the conserved kinase and hub domains and high variability in the linker region makes it difficult to specifically identify splice variants of the CaMKII paralogs in cellular experiments.

**Figure 2.**
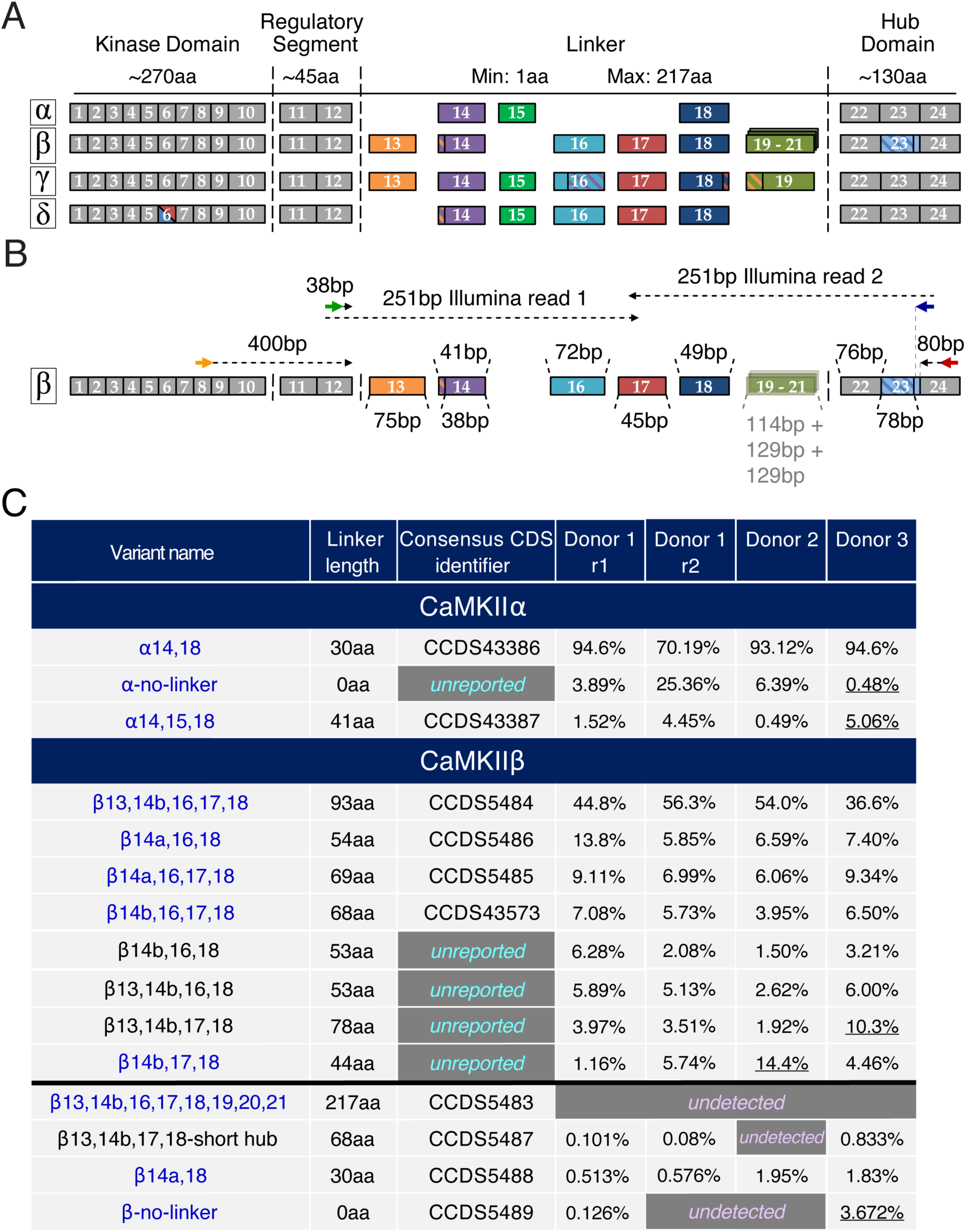
Illumina sequencing reveals a heterogeneous population of CaMKII variants in the human hippocampus. (**A**) Human CaMKII genes are encoded by 24 exons with clear one-to-one correspondence across genes. Exons 1-10 encode the catalytic kinase domain, 11 and 12 encode the regulatory segment, 13-21 encode the variable-length linker, and 22-24 encode the association (hub) domain. Missing linker exons indicate that the exon is not encoded in the corresponding gene. Constitutively incorporated kinase, regulatory segment, and hub exons (grey) are spliced into every transcript. CaMKIIδ encodes two versions of kinase exon 6, with one or the other, but not both, incorporated into transcripts. Several exons contain alternative splice sites, permitting omission of a fraction of the exon (indicated by hatching). CaMKIIβ encodes three nearly identical proline-rich linker exons: 19-21, homologous to CaMKIIγ exon 19. (**B**) Illumina library construction is illustrated with CaMKIIβ. Initial PCR amplification of a region encompassing all alternative splice sites (including the linker and hub exon 23) is carried out with primers designed for specificity for target CaMKII gene, to the exclusion of the other three genes (orange and red arrows). Initial amplicons serve as templates for amplification of library inserts in a second PCR reaction with a different primer pair (green and blue arrows). Resulting inserts are fully spanned by paired-end 251bp reads for nearly all splice variants. For CaMKIIβ, this is true as long as exons 19, 20, and 21 are not incorporated. These exons are shown as semi-transparent because they were not incorporated in any detected hippocampal splice variant. (**C**) Summary of CaMKIIα and β transcript variants for 2 replicates of Donor 1 (r1, r2), Donor 2, and Donor 3 as detected by Illumina sequencing. All detected variants, including all previously annotated variants, are shown for CaMKIIα. For CaMKIIβ, the top eight read-mapping variants are shown above the black line and the remaining previously annotated variants are shown below the line. Variants are marked as ‘unreported’ if they have not been previously annotated or ‘undetected’ if we did not detect the transcript in our experiment. Underlined percentages indicate inconsistency between libraries. Variants highlighted in blue are those that were used in activity measurements (see Fig. 3).

There have been many studies that outline specific roles for exons in the variable linker region, which provide some clues as to why exons may be spliced in or out. Exon 15, which is encoded in all genes with the exception of CaMKIIβ has been documented to contain a nuclear localization sequence (NLS) (*20–22*). This NLS (KKRK) is a common motif identified as a sufficient sequence to achieve nuclear localization (*23*). Exon 13, which is encoded in CaMKIIβ and γ, but not α or δ, mediates an important interaction with F-actin (*24, 25*). In dendritic spines, CaMKII interacting with actin is crucial for the maintenance of spine structure during long-term potentiation (*26*). Perhaps the most dramatic form of alternative splicing results in the formation of alpha CaMKII associated protein (alpha KAP) where CaMKIIα is spliced to a form that includes only the hub domain (*27*). This splice variant plays an important role in membrane anchoring using its hydrophobic N-terminal domain (*28*). Finally, it was shown that alternative splicing within the hub might regulate oligomerization (*5*). Roles for the remaining exons have yet to be delineated.

In addition to roles assigned to specific exons, the variable linker region has also been shown to play a critical role in CaMKII activation. A standard metric for CaMKII activity is the concentration of Ca^2+^/CaM required for half-maximal activation (EC_50_ value) (*19, 29*). It has also been clearly shown that CaMKII activation is cooperative (*29*). In regard to linker length, a CaMKIIα variant with a 30-residue linker has been shown to require substantially less Ca^2+^/CaM to achieve maximal activity compared to a CaMKIIα variant with a 0-residue linker (*19*). This general trend also holds in comparing a long linker CaMKIIβ variant to a short linker CaMKIIα variant (*30*).

Here, we address the gaps in knowledge in three major areas: 1) which specific CaMKII splice variants are expressed in human hippocampal cells, 2) how splicing affects CaMKII activation, and 3) structural analysis of an autoinhibited CaMKII holoenzyme with a 30-residue linker.

## Results

### The exon architecture is conserved between CaMKII paralogs

The four genes encoding CaMKII in humans share a high degree of homology. Using the Neighbor Joining method (*31, 32*), we created reconstructions of the kinase and hub domain divergence for the four human CaMKII genes rooted with D. melanogaster CaMKII kinase and hub outgroups (Fig. 1D). Because of the clear one-to-one correspondence of exons across genes, we employed a common exon numbering system (Fig. 2A). Herein, we implement a new naming scheme for CaMKII variants based on ^1^gene identity and ^2^incorporated exons. Moving forward, we propose this is the simplest way to efficiently catalog current CaMKII variants and include new variants as they are discovered.

### Deep sequencing revaels at least 70 CaMKII transcript variants in human hippocampus

We carried out targeted deep sequencing of hippocampal CaMKII transcripts from three separate human donors (82, 26, and 66 year-old males) without neurological or neuropsychiatric diagnoses. In order to identify all alternatively spliced CaMKII transcripts expressed in a human total RNA sample, these transcripts must be substantially enriched. Similar to earlier approaches (*1,3,33-35*), we reverse-transcribed human hippocampal RNA to generate cDNA and then used gene-specific primer pairs to PCR-amplify variable regions of CaMKII transcripts (Fig. 2B).

Agarose gel separation of PCR amplicons followed by Sanger sequencing of excised gel bands was inadequate for detailed characterization of alternative CaMKII splicing, because many bands contained mixtures of variants (fig. S1). To circumvent this issue, we devised an Illumina next-generation sequencing approach. Given the locations at which CaMKII transcripts are variably spliced and the maximal lengths of variable regions when all optional exons are incorporated, strategic placement of a primer pair facilitates PCR-based construction of Illumina sequencing libraries such that 251bp paired-end sequencing allowed unambiguous identification of virtually all possible CaMKII transcripts from all four genes (Fig. 2B, blue/green primer pair). Compared to traditional whole-transcriptome Illumina RNA-Seq library preparation, this approach emphasizes sensitivity of unambiguous variant detection over accuracy in quantification of relative transcript abundance, due to the greater potential for amplification bias resulting from additional PCR cycles. Therefore, we make no claims with respect to transcript abundance beyond classifying transcripts into several detection level categories based on the number of reads mapped to each transcript. However, spurious detection of transcripts is extremely unlikely, since we applied a stringent similarity cutoff of a minimum of 10 reads mapping to each transcript (see Materials and Methods).

We performed this experiment with three separate donor samples of human hippocampal tissue. We detected a surprisingly large number of distinct transcript variants (Fig. 2C, Tables S1-S16, data file S1), including 3 CaMKIIα variants and over 15 each of CaMKIIβ, CaMKIIγ, and CaMKIIδ. In addition to variants previously annotated in genomic databases (NCBI Consensus CDS collection, NCBI GenBank, Ensembl), we also identified numerous previously unobserved variants of each gene, which we refer to as ‘unreported’ in this manuscript. One unreported variant of note was the CaMKIIα transcript including no optional linker exons (0 aa linker). This is significant because the crystal structure of CaMKIIα no-linker variant is the only available high resolution structure of full length CaMKII (pdb: 3SOA (*19*)), but this variant was previously not known to be biologically relevant.

Nearly every identified transcript incorporating optional linker exons contained exons 14 and 18, suggesting these two exons constitute a “core” linker predating the divergence of the four genes. Assuming the core is mandatory if any other linker exons are to be spliced in, we observed a large fraction of allowable linker sequences for each gene. CaMKIIβ variants incorporating proline-rich exons 19, 20, or 21 and CaMKIIγ variants incorporating F-actin-interacting exon 13, which were not detected in our experiment, are two notable exceptions.

We assessed the reproducibility of this experiment with two replicates of sample preparation and sequencing for Donor 1, the highest quality tissue and RNA sample (Fig. 2C, Tables S1-S4). For these technical replicates, 99% of reads mapped to the same collection of variants. Not surprisingly, variants on the edge of detection (cumulatively mapping to 1% of reads) differed between the replicates (Tables S2, S6, S10, and S14). However, there was too much variability between the replicates to precisely quantify detection levels of variants and confidently order them based on detection level (Materials and Methods, Tables S17-S19). Instead we bin variants into several detection-level categories.

Expressed variants were also quite consistent across donors, with at least 97.5% of all reads from each donor mapped to a collection of variants detected in all three donors (Tables S1-S16). Most, but not all, variants fell into the same broad expression category across all donors. Accordingly, we designated variants as ‘consistent’ and ‘inconsistent’ (Tables S1, S5, S9, S13). Large differences in detection levels between donors observed for inconsistent variants may reflect underlying physiological differences between donors.

### The variable linker affects activation of CaMKIIα, but not CaMKIIβ

We focused on the activation properties of CaMKIIα and β variants because of their well-documented involvement in learning and memory (*9, 11*). First, we determined the EC_50_ values for Ca^2+^/CaM activation of all three CaMKIIα variants sequenced from human hippocampus. For all EC_50_ measurements, the curve fit is shown for simplicity (Fig. 3), and fits with all data points used are shown (fig. S2). For brevity we refer to variants by linker length: α-41 (α14,15,18 in Fig. 2C), α-30 (α14,18), and α-0 (α-no-linker). In agreement with previous experiments comparing α-30 to α-0 (*19*), we also observed a large right-shift in the EC_50_ value (from 24 nM to 313 nM, respectively, Fig. 3A). CaMKIIα-41 (EC_50_ = 25 nM, Fig. 3A) did not show a significant difference compared to α-30.

**Figure 3.**
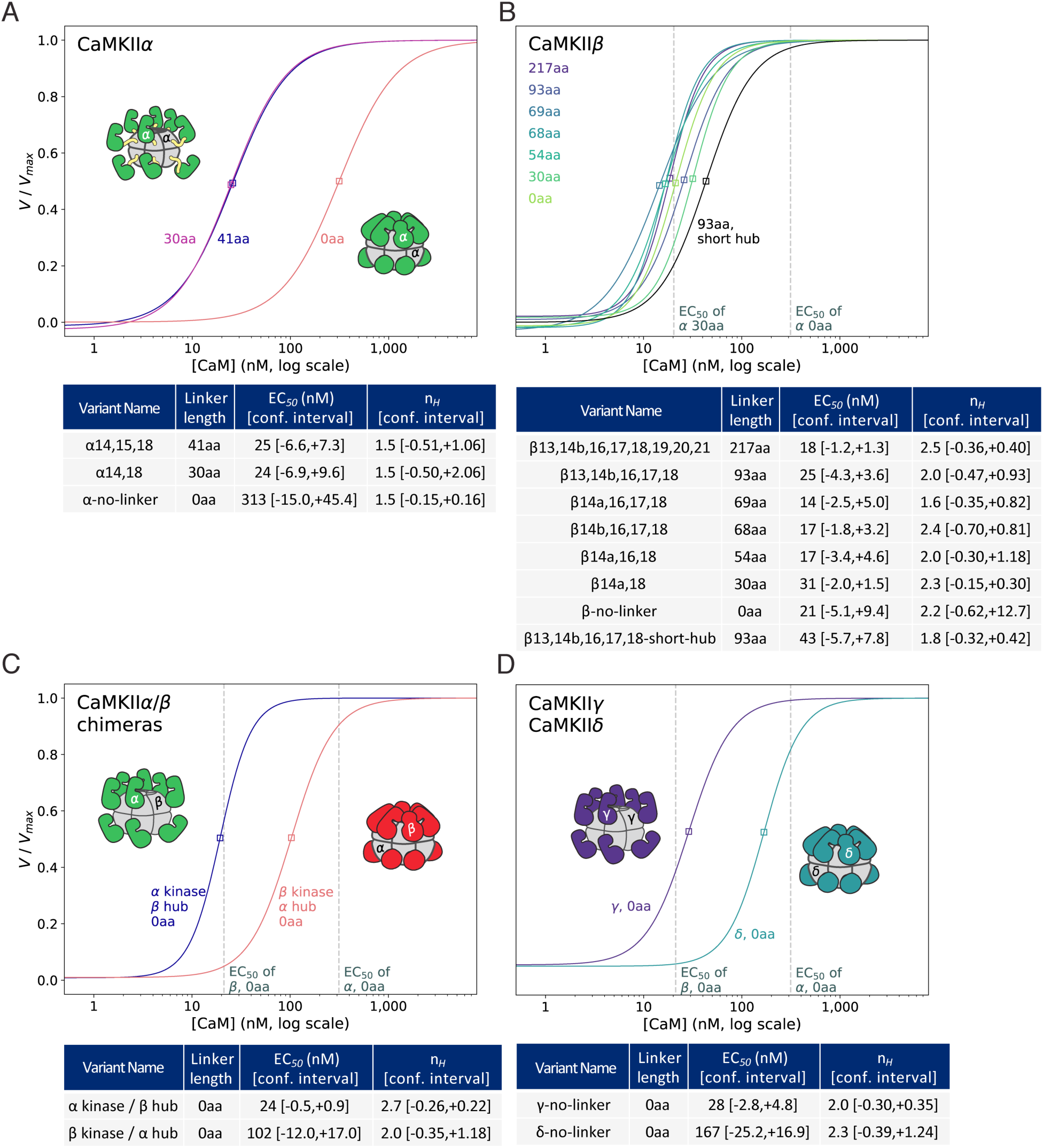
The hub domain plays a crucial role in Ca^2+^/CaM sensitivity. CaMKII activity against a peptide substrate (syntide) was measured as a function of Ca^2+^-CaM concentration. To simplify comparisons, fit curves were normalized by the corresponding V_max_ fit parameter and data points are not shown (see fig. S3). The EC_50_ value for each fit is indicated by a single box, and dashed gray lines in B-D indicate reference EC_50_ values for CaMKIIα-30 and CaMKIIα-0 (B) or CaMKIIβ-0 and CaMKIIα-0 (C, D), as labeled on the X-axis. Activity measurements and EC_50_ and n_H_ (Hill coefficient) fit parameters are shown for all three CaMKIIα splice variants (**A**), eight selected CaMKIIβ variants (**B**), chimeras of CaMKIIα/β with no linker (**C**), and CaMKIIγ and δ with no linker (**D**). EC_50_ and Hill coefficient (n_H_) fit parameters with 95% confidence intervals (see Materials and Methods) are listed for each tested variant, N=3 titrations for each protein sample. Cartoon representations of CaMKII holoenzymes in their autoinhibited states are shown in panel A. For panels C and D, each domain is labeled according to the CaMKII gene.

Next, we examined CaMKIIβ variants with the broadest possible range of variable linker lengths: previously annotated and detected variants β-93, β-69, β-68, β-54, β-30, and β-0, as well as the annotated, but undetected, variant β-217 (Fig. 3B). Quite surprisingly, we observed no major differences in EC_50_ across the entire range of linker lengths, with EC_50_ of all β variants falling in the narrow range between 14-31 nM. This result showed that the variable linker does not tune activation the same way in different CaMKII genes. The Hill coefficients (n_H_) from these measurements of CaMKIIα and β variants were all quite similar, ranging from 1.5 to 2.5, which was consistent with previous measurements (*19*). Of note, the cooperativity in CaMKIIβ variants were consistently higher compared to CaMKIIα, but did not correlate with EC_50_ value differences.

We observed a 15-fold difference in the EC_50_ for Ca^2+^/CaM between α-0 and β-0 (313 nM vs 21 nM). The difference in activation properties of these two variants may be attributed to sequence differences outside of the variable linker: in the kinase domain, the hub domain, or a combination of the two. In fact, the only CaMKIIβ variant we tested with an EC_50_ outside of the narrow range around 20 nM was the novel variant β-93-short-hub, which is slightly right-shifted from the other CaMKIIβ variants (Fig. 3B). The only difference between β-93 (EC_50_ = 25 nM) and β-93-short-hub (EC_50_ = 43 nM) are 26 residues missing in the hub domain, indicating that hub alone may affect activation properties. These results motivated our next experiments.

### Hub identity plays a regulatory role in holoenzyme activation

We directly assessed the regulatory contribution of the kinase and hub domains of CaMKIIα and β (90% and 77% identity, respectively). To do this, we created two chimeras with 0aa linkers. The first chimera was the CaMKIIα kinase domain and regulatory segment fused to the CaMKIIβ hub domain, and the second chimera was the CaMKIIβ kinase domain and regulatory segment fused to the CaMKIIα hub domain. The EC_50_ value of the β kinase/α hub chimera was substantially right-shifted compared to that of the α kinase/β hub chimera (Fig. 3C). Both CaMKIIβ-0 and the chimera with the β hub had roughly the same EC_50_ (24 nM and 21 nM, respectively). On the other hand, CaMKIIα-0 and the chimera with the α hub were both right-shifted. However, the EC_50_ of the chimera was still 3-fold lower than the EC_50_ of wild type CaMKIIα-0: 102 nM vs 313 nM. This suggested that the role of the hub domain in determining activation properties was dominant with respect to the kinase domain, with no contribution from the kinase in the context of β hub, but a significant contribution in the context of α hub.

### Activation properties track with evolutionary divergence

We compared the divergence of human CaMKII kinase and hub domains using the corresponding domains from the single CaMKII gene in fly as outgroup to root the trees (Fig. 1D). The kinase and hub trees have consistent topologies, in which the common ancestor of CaMKIIα and δ diverged from the common ancestor of β and γ. However, while the four kinase domains diverged in nearly a star topology (the segments separating the α/δ and β/γ ancestors from their common ancestor are much shorter than any other segment), hub divergence was not nearly as uniform. CaMKIIα and δ are much more diverged from each other and from β/γ.

CaMKIIβ and γ hubs have diverged by far the least, with the fewest substitutions between them. Based on this similarity, and the apparent involvement of the hub domain in activation, we hypothesized that γ-0 would behave similarly to β-0. We measured activation of γ-0 and δ-0 (Fig. 3D), confirming our hypothesis (γ-0: EC_50_ = 28 nM). Consistent with the order in which α, β/γ, and δ split, as well as with the greater degree of divergence between α and δ hubs, δ-0 required an intermediate amount of Ca^2+^/CaM for activation (EC_50_ = 167 nM): roughly 7-fold higher than β-0, but 2-fold lower than α-0.

### Insight into the structural mechanism of activity regulation

The striking difference we observed in the activation of CaMKIIα-0 compared to CaMKIIβ-0 (EC_50_ = 313 nM and 21 nM, respectively) led us to investigate whether there was a structural explanation. The crystal structure of CaMKIIα-0 shows a compact conformation with a radius of gyration (R_g_) of 61 Å as determined using small angle X-ray scattering (SAXS) (19). We performed SAXS measurements to compare the overall shape of CaMKIIα-0 and CaMKIIβ-0 in analyses revealed a significant difference in the R_g_ and D_max_ of CaMKIIα-0 (∼59 Å and 155 Å, in line with previous data (19)) compared to CaMKIIβ-0 (∼66.4 Å and 250 Å) (fig S3). These results showed that CaMKIIα-0 is in a more compact conformation than CaMKIIβ-0. Our current understanding of the CaMKII holoenzyme state (compact versus extended) has emerged from a combination of negative stain EM, SAXS, and X-ray crystallography, as reviewed by (31). The central hub is consistently observed as a dodecameric or tetradecameric assembly. The kinase domains have been observed in multiple conformations, including above and below the hub domain plane in EM studies (32), in compact positions docked against the hub in an X-ray crystal structure (19), or in the same plane as the central hub at extended positions, from EM and SAXS modeling (33, 34). Another negative stain EM study of a CaMKIIα-30 construct showed the kinases adopting positions at extended distances from the central hub (35).

A general challenge for 3D EM reconstructions of CaMKII has been the strong preferred orientation of the sample under both cryo and negative stain conditions. In addition, the average size of negative stain particles and the linker distances present in CaMKII isoforms have motivated careful comparison of negative stain and cryo-conditions when interpreting kinase domain position (32).

To revisit questions surrounding the CaMKII holoenzyme conformational state, we analyzed samples of CaMKIIα-30 using single particle cryo-EM. 2D class averages of the sample, frozen at 0.5-1 mg/ml, showed clearly distinguishable 6-fold and 7-fold versions of the central hub with visible secondary structure elements (Fig. 5A). Obvious side views were visible in the 2D classes, in contrast with previous negative stain analyses (Fig. 5B) (17,35,36). 2D classes also showed additional, less ordered density, proximal of the hub (presumably from the kinase domains) (Fig. 5C, D). This density was still present when the mask diameter was expanded during 2D classification (Fig. 5E). No additional ordered density, at more extended positions, appeared to be excluded due to masking. In top views of 6- and 7-fold hub assemblies, additional kinase domain density was observed as a strong signal positioned between hub subunits (Fig. 5C), and was observed in side-views above and below the mid plane of the hub (Fig. 5D, E).

We generated initial models using Relion initial model, and classified the full particle set, all without any assumptions of symmetry (see Materials and Methods for details). Both 6-fold and 7-fold hub domains (without any ordered kinase domains) emerged during 3D classification.Ordered density of the kinase domains appeared in two final classes. The first class, which we refer to as Class A, was refined to 4.8 Å resolution (as evaluated by gold standard FSC (Fourier Shell Correlation) criterion). In Class A we observed one ordered kinase domain, docked against a 6-fold hub in a position with the beta-clip engaged with the hub, as seen in the crystal structure of CaMKIIα-0 (pdb: 3SOA) (Fig. 5). A second class (Class B) was refined to 6.6 Å resolution. Class B showed one kinase domain interacting with a 6-fold hub in a new orientation. In Class B, the kinase C-lobe (helices: αEF and αG) engaged a loop in the hub domain (W373-P379), and the N-lobe was in a position closer to the hub mid-plane. In both reconstructions, the electron density (with recognizable secondary structure elements) was unambiguous for positioning the kinase domain. Both Class A and B exhibited preference for top views, however these data showed angular distributions which allowed 3D reconstruction (fig. S4). Similar docked states as Class A and B were seen with 7-fold hubs. Enforcing D6 symmetry during refinement of the Class A or Class B particles resulted in reconstructions with a recognizable central hub, but loss of recognizable kinase density.

## Discussion

We unambiguously identified at least 70 distinctly spliced CaMKII transcripts present in three individual human hippocampal samples using Illumina sequencing. It is important to note that excised hippocampal tissue samples are necessarily comprised of multiple cell types from the hippocampus, including neurons, glia, interneurons, etc. Therefore, the transcripts we report here are derived from this mixture of cell types, not from excitatory neurons alone. From our experiment, it is impossible to tell which transcripts are co-expressed specifically in excitatory neurons or any other cell type. Addressing this important question requires precise extraction of individual neurons from the hippocampus followed by single-cell transcript sequencing. The alternative, culturing neurons outside of brain tissue, disrupts numerous *in vivo* cell-cell interactions, including synapses, quite possibly leading to non-physiological changes in CaMKII transcription and splicing.

Using the sequences we obtained, and others previously annotated from various tissues, we examined the differences in kinase activity regulation between CaMKII genes and splice variants. We discovered that, while CaMKIIα regulation is dependent on the variable linker region, CaMKIIβ regulation is not. Further, we now show a role for the hub domain in the regulation of CaMKII activation by Ca^2+^/CaM. Our data provide important insights into the heterogeneity of CaMKII transcripts in a human tissue and uncover a new role for the hub domain in regulating CaMKII activity through an allosteric mechanism. This finding opens a new window to allosteric control of CaMKII activity through modulation of the hub domain.

We sequenced a large number of transcripts from all four CaMKII genes from 3 separate samples of human hippocampal tissue. Notably, CaMKIIα produced a no-linker transcript (CaMKIIα-0) that accounted for 0.5% - 25% of mapped reads. This sequence was previously crystallized to obtain the only full-length atomic resolution structure of CaMKII. Our data demonstrate the potential physiological relevance of the compact conformation observed in that structure (*19*). The sheer number of detected variants detected in the hippocampus (70 across the four CaMKII genes) suggests that a) either splicing is not tightly regulated at every exon, or b) there are complex mechanisms of regulation responsible for every transcript we detected. In either case, certain exons (proline-rich CaMKIIβ exons 19-21 and F-actin-interacting CaMKIIγ exon 13) appear to be specifically suppressed in the hippocampus.

In terms of abundance of detected transcripts, systematic PCR bias is unlikely to be solely responsible for the roughly 1000-fold differences between the high- and trace-level detection categories of detected variants of CaMKIIβ, γ, and δ (Tables S1, S5, S9, S13). Physiological differences in transcript copy number between variants likely contributed to these differences in detection level. Furthermore, given the overall consistency in variant detection between donors, the few instances of dramatic detection level differences may reflect genuine physiological differences between individuals.

We found that agarose gel separation of transcripts followed by Sanger sequencing of individual bands cannot resolve CaMKII variants from hippocampus, because sequencing traces from most bands clearly indicated a mixture of templates present in the sequenced sample. In many cases, careful analysis of the Sanger sequencing trace allowed us to identify at least two distinct splice variants (fig. S1). Therefore, it is difficult to interpret the presence or absence of an expected gel band as indicative of the presence or absence of a particular CaMKII transcript in a sample.

Since each Illumina read obtained by our approach reflects a single template molecule, it produces a more robust survey of the CaMKII transcripts. We suggest that this approach should be applied to other tissues to begin to catalog the diversity in CaMKII transcripts. We expect that in other tissues the identity and relative abundance of transcripts will vary from those we observed in the hippocampus. With this information in hand, we will certainly improve our understanding of the regulation of CaMKII alternative splicing, and ultimately be able to link this regulation to the Ca^2+^ signaling requirements within each tissue.

Our results comparing the EC_50_ values between CaMKIIα and β splice variants show that the regulation of CaMKII activation is gene-dependent and, specifically, affected by the sequence of hub domain. Our measurements characterize CaMKII activation at equilibrium Ca^2+^/CaM concentrations. CaMKII activity is also dependent on the frequency of Ca^2+^ spikes, which is how Ca^2+^ is delivered to endogenous CaMKII in post-synaptic neuron (19,30,37). Future experiments will be needed to carefully dissect this kinetic contribution in activation of different splice variants. The variable linker in CaMKIIα does play a role in regulating CaMKIIα activity, which is consistent with previous work (*19*). We used cartoon holoenzymes illustrate the compact and extended autoinhibited states of CaMKII (Fig. 3), which correspond to lower and higher sensitivities to Ca^2+^/CaM, respectively. For CaMKIIα, these cartoons are based on a crystal structure (pdb: 3SOA) and SAXS measurements, respectively (*19*). Surprisingly, the variable linker in CaMKIIβ does not play a role in regulating CaMKIIβ activity. Our SAXS measurements directly comparing CaMKIIα-0 and β-0 suggest that α-0 adopts a compact conformation whereas β-0 adopts a more extended conformation as evidenced by its larger R_g_ and D_max_ (fig. S3). This data corroborates the trend that was previously observed comparing α-0 which is what we observe in our experiments as well.

Our results with CaMKIIα and β chimeras provide evidence that the hub identity is a determinant of sensitivity to Ca^2+^/CaM. We observed a large difference in activation of the CaMKIIβ kinase when it was fused to the CaMKIIα hub domain. Substitutions between CaMKIIα and β hub domain sequences are distributed quite uniformly throughout the domain (Fig. 4A). This led us to hypothesize that there is a dynamic component to this regulation. We turned to single particle cryo-EM to gain a better understanding of the structural mechanism of activity regulation.

**Figure 4.**
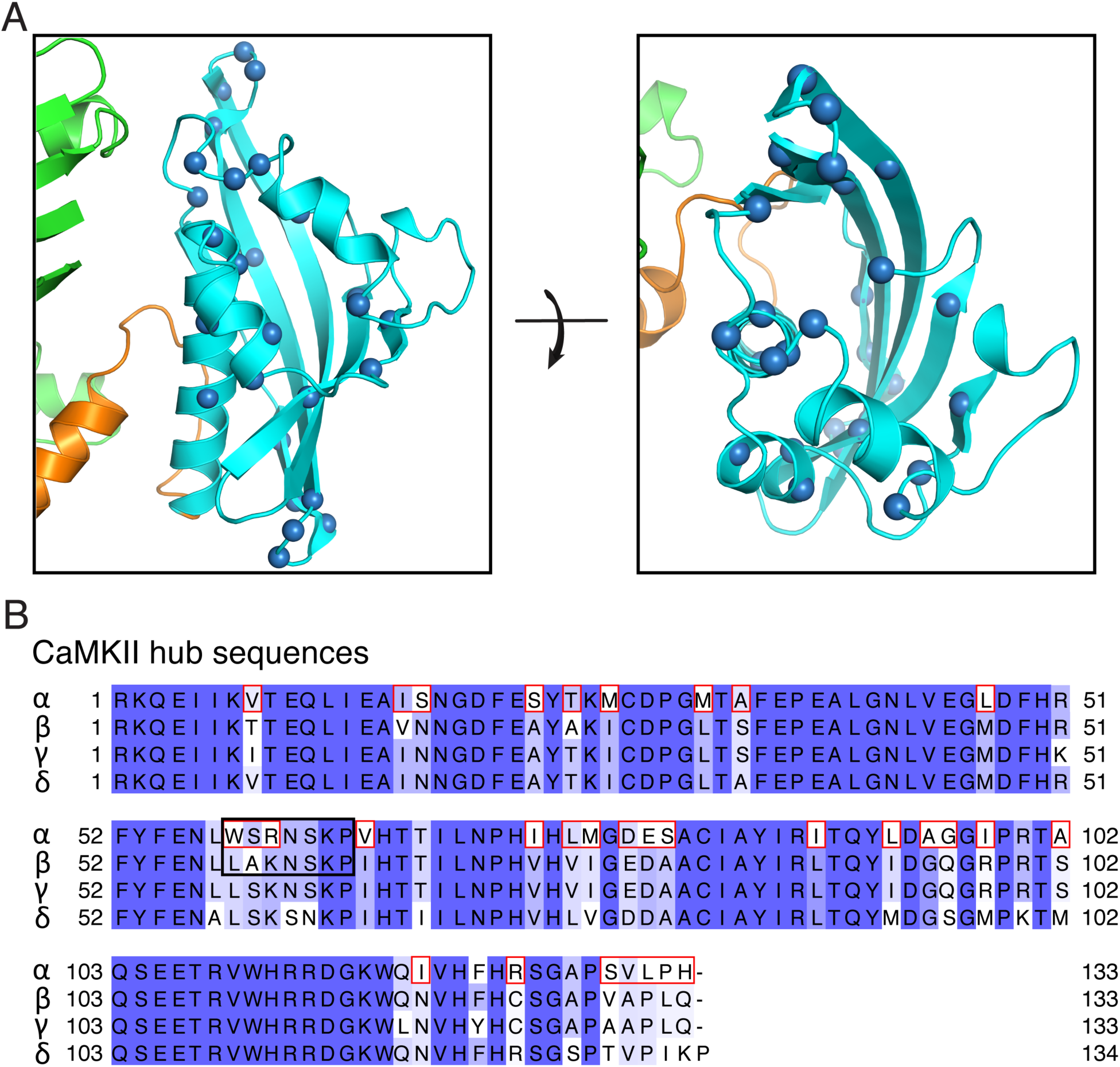
Allosteric contributions from the hub domain regulate CaMKII activation. (**A**) The 32 positions that differ between CaMKIIα and β hubs are represented by spheres at the alpha Carbon position on the structure of CaMKIIα hub (pdb: 3SOA). Hub is shown in cyan, regulatory segment in orange, and kinase domain in green. (**B**) Sequence alignment of CaMKIIα, β, γ, and δ hub domains, colored by degree of conservation (dark to light = most to least conserved). For convenience, all hub sequences start at residue 1. Residues in CaMKIIα hub that differ in CaMKIIβ are highlighted by red boxes. The loop mediating kinase docking in the Class B cryo-EM reconstruction is shown in the black box.

Our cryo-EM data show that the kinase domains are in close physical contact with the hub, and suggest two specific sources for hub-dependent regulation observed in the chimera experiments. These cryo-EM data clearly observe two conformations sampled by the kinase domain under frozen hydrated conditions. First, we observe the same docked conformation in single-particle cryo-EM reconstructions using CaMKIIα-30 (Fig. 6A, Class A) that was observed in a crystal structure of CaMKIIα-0 (19). The crystal structure shows a compact state of the holoenzyme where the kinase domain is docked onto the hub domain and the calmodulin binding segment is occluded. The conformation we observe in cryo-EM (Class A) suggests that with the holoenzyme in solution, kinase domains may sample a similar compact state, even in the presence of a 30-residue linker. Single amino acid substitutions from CaMKIIα to β are not found at this docking interface, thereby confounding straightforward tests of this interface in hub-dependent regulation. Our cryo-EM analysis also revealed a new second, distinct docked state, Class B (Fig. 6B). In this reconstruction, the kinase domain is flipped relative to Class A, with the C-lobe oriented above the N-lobe, relative the hub. A loop in the hub domain mediating interaction is variable between CaMKIIα and β (Fig. 4B, black box). In Class B, the regulatory segment is accessible for calmodulin binding, but there remains extensive contact between the kinase C-lobe and the hub, which may influence activation. These two cryo-EM reconstructions advance our understanding of holoenzyme architecture because they show that in a holoenzyme with a 30-residue linker the kinase domains make intimate contact with the hub, to the extent that two distinct cryo-EM reconstructions can be observed with specific docked conformations.

We note that in these reconstructions, only one kinase domain was sufficiently ordered to be resolved in the final reconstruction of a dodecamer. The final reconstructions for Class A and B were obtained with masks that excluded non-modeled density, as standard. Outside of this masked region was unaccounted density present above and below the midplane, as observed in the 2D class averages (Fig. 5). This significant disordered density may reflect extensive transient interactions between the hub and kinase domains. The position of the unmodeled density is consistent with kinase domain positions along the surface of the hub, and incompatible with the extended positions proposed in the negative stain reconstructions (35). It will be interesting to investigate whether constructions like CaMKIIα kinase/β hub chimera show this kinase proximity to the hub or not. The 30-residue linker in CaMKIIα-30 is predicted to be a disordered segment. This disordered linker allows the kinase domains many degrees of freedom. Locking the kinase into a docked conformation is likely to be entropically unfavorable and transient, making coincident docking statistically unlikely.

**Figure 5.**
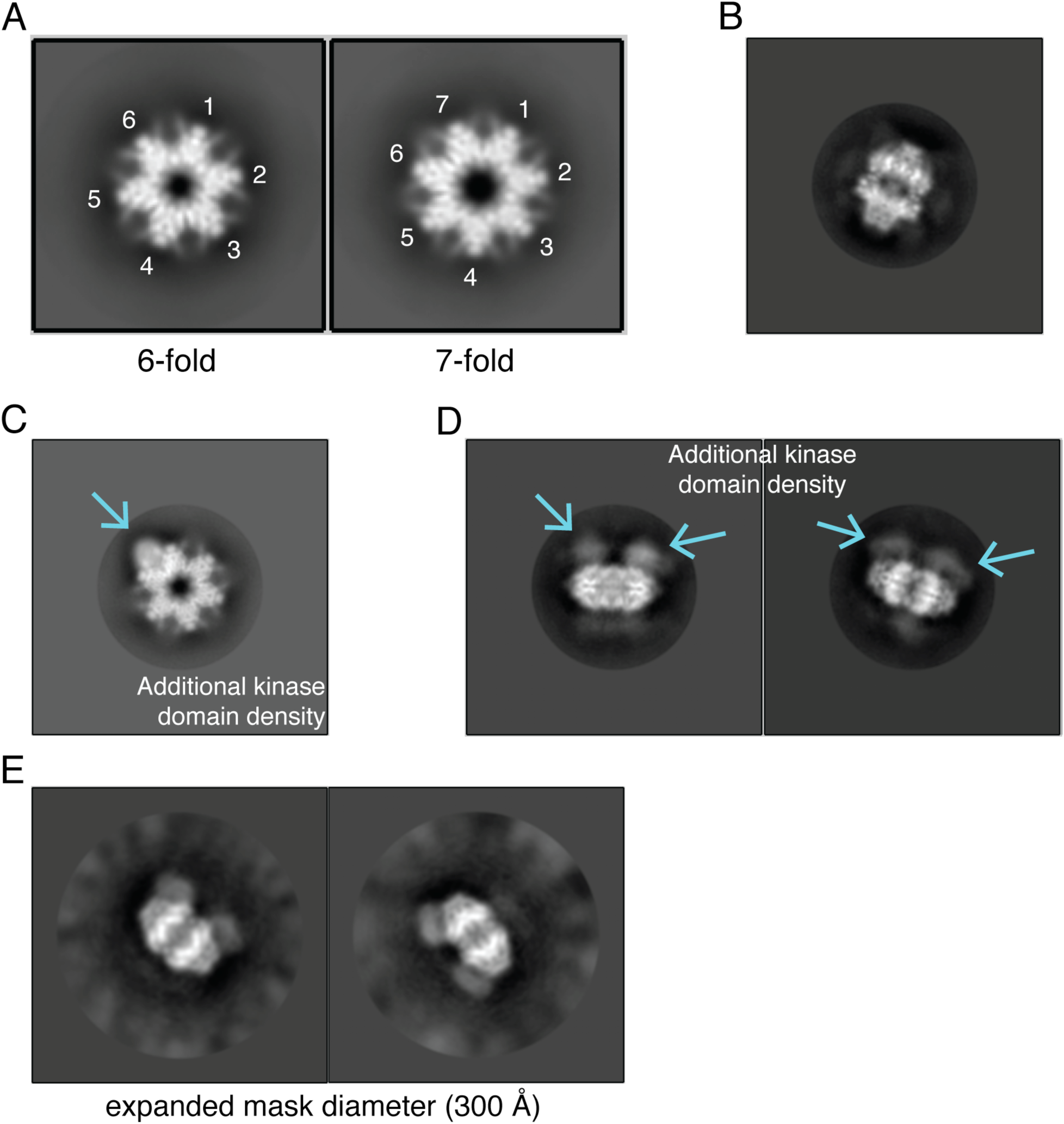
2D classes of single particle cryo-EM analysis of CaMKIIα-30. (A**) 6- and 7-fold** symmetric top views for CaMKII holoenzymes were observed cryo-EM 2D classes. (B) Example of a clearly visible side views in the 2D classes. (C) Density for a single kinase domain (blue arrow) docked onto a dodecameric central hub seen in a top view. (D) Density for multiple kinase domains (blue arrows) above the hub midplace is visible in side views. (E) Density for kinase domains remain when the mask diameter is increased to 300 Å. No additional ordered density is visible at extended positions.

Our cryo-EM analysis shows two 3D reconstructions of CaMKIIα-30 with different docked states in which the kinase domain makes direct contact with the hub. We speculate that these interactions may mediate autoinhibition or calmodulin-dependent activation. As the interactions we observe may vary depending on the identity of the CaMKII hub domain, they do not exclude other interactions, which may be uncovered when kinase domains are less likely to interact with the hub (34). With the rich diversity of splice variants described here, a given holoenzyme may have an entire set of regulatory interactions it can sample. Thoughtful future experiments will be needed to specifically distinguish the role of these dynamic interactions in holoenzyme activation, to more fully elucidate the intricate mechanistic details of a remarkably versatile enzyme.

**Figure 6.**
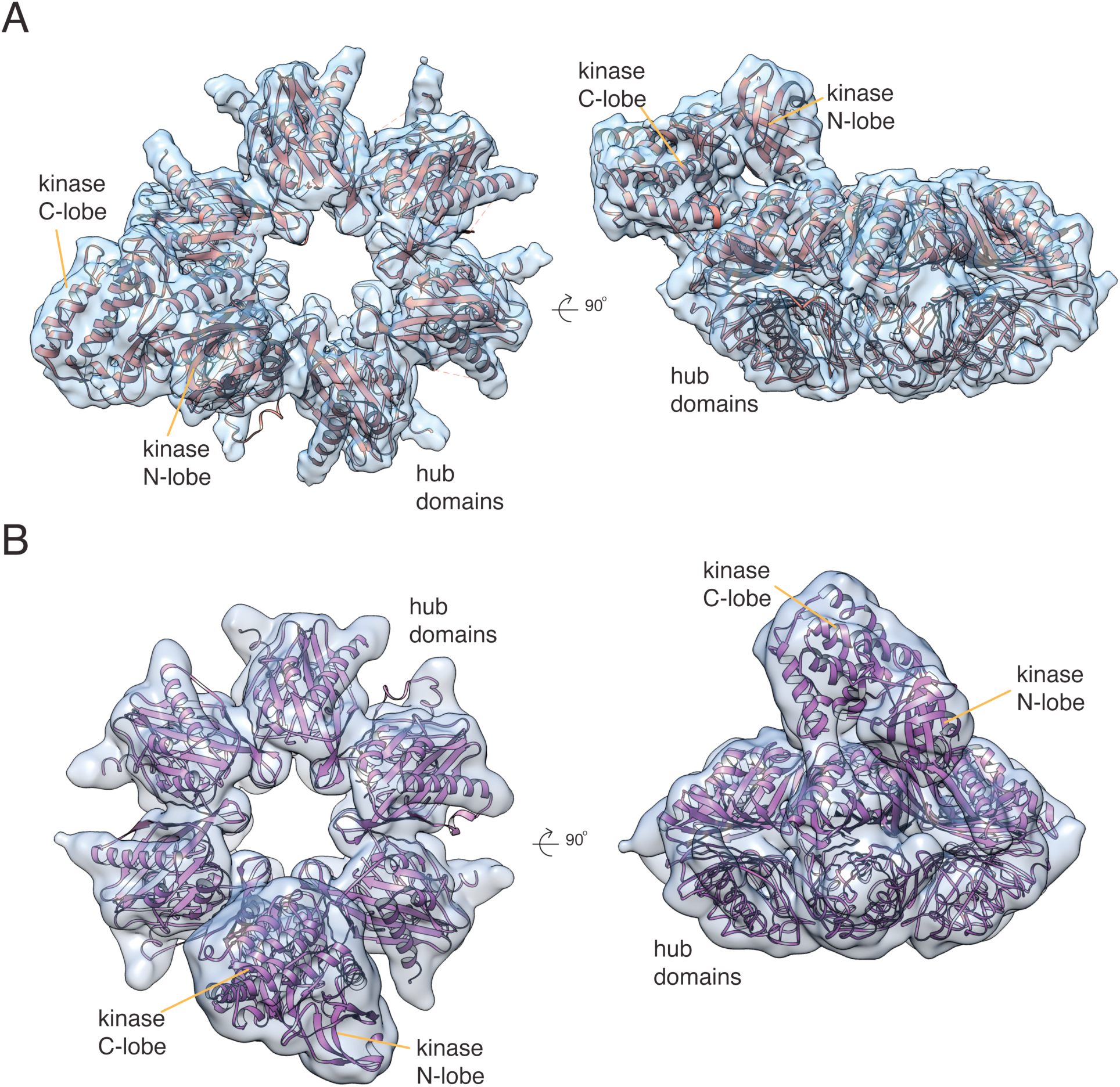
Two kinase domain arrangements observed by cryo-EM. (A) In reconstruction Class A, a single kinase domain is resolved, between two hub domain subunits. A kinase domain from the CaMKII**α**-0 holoenzyme crystal structure (pdb: 3SOA) was fit into the additional density. (B) In a second reconstruction, Class B, a kinase domain is visible interacting with a hub subunit with the kinase C-lobe making contact with a loop at the top of the hub domain. CaMKII**α** kinase domain (pdb: 3SOA) docked into the additional density, CaMKII hub domain (pdb: 5IG3) docked into the hub density.

## Materials and Methods

### Initial amplification of transcript variable regions

Two human total RNA samples and one human cDNA sample were commercially purchased from BioChain. The samples were derived from complete hippocampi of three human donors (donor 1: 82 year old male, donor 2: 26 year old male, donor 3: 66 year old male) without neurological or psychiatric diagnoses. BioChain advised that the tissue from donor 2 did not pass all internal histological quality controls, although the RNA derived from that tissue did pass all RNA quality controls. We detected many fewer low-expression variants (less than 1% of mapped reads) of CaMKIIβ, γ, and δ in the sample from donor 2, compared to donors 1 and 3. However, robustly expressed variants were highly consistent between all three donors (Tables S1-S16).

This is consistent with non-specific partial degradation of RNA in the donor 2 sample. While low copy number transcripts fell below the threshold of detection due to partial degradation across all mRNAs, robustly expressed transcripts remained detectable, despite losing a fraction of their initial copy number.

cDNA was reverse-transcribed from donor 1 and donor 2 RNA samples using the ProtoScript II first strand cDNA synthesis kit (NEB #E6560) with oligo dT priming. Variable regions of transcripts were PCR-amplified for 35 cycles using high-specificity primer pairs designed to minimize cross-hybridization between CaMKII genes (Supplementary table XX) using either Phusion polymerase (NEB #M0530) or KAPA HiFi polymerase (Roche). This yielded a mixed population of variable-length amplicons reflecting the splice variants of the targeted gene present in the starting sample.

### Construction of Illumina Sequencing Libraries

Illumina sequencing libraries were prepared from the initial variable-length amplicon pools using an adapted PCR-based construction protocol (38). Briefly: first, gene-specific primers with overhangs containing Illumina-specified sequences (Fig. 2B, supplementary file S2) were used to sub-amplify transcript variable regions. Second, generic primers annealing to the overhangs introduced in the first reaction were used to incorporate annealing sites for Illumina sequencing primers, NEBNext library indices (NEB #E7335 single-index set, #E7600 dual-index set), and P5/P7 flow cell annealing sequences. The resulting amplicons were agarose gel-purified with size selection based on the theoretical minimal (no linker exons spliced in) and maximal (all linker exons spliced in) lengths of the variable region inserts.

### Analysis of Illumina Sequencing

Illumina reads were grouped by sequence similarity, producing clusters with >90% sequence identity and two or fewer base pair insertions/deletions. The consensus sequence of each cluster was aligned to exon sequences from the corresponding CaMKII gene obtained from the reference human genome (GRCh38.p12, Ensembl release 95) in order to map exon order in the transcript represented by the cluster. Over 90% of clustered reads from each Illumina library were unambiguously mapped with reference exons of the corresponding gene, with all splice junctions matching those observed in previously annotated splice variants. We refer to these as mapped reads. Python2 code implementing read clustering is available at github.com/romansloutsky/illumina_read_clustering.

Two technical replicates of libraries were prepared from the donor 1 tissue sample, starting with the initial reverse transcription step, and sequenced. One replicate each from donors 2 and 3 was sequenced. For each gene, different collections of splice variants were detected in each of the four cDNA samples, so calculating the correlation between replicates across all detected variants is not possible. Instead, for each gene we calculated pairwise Pearson correlation in variant detection levels (fraction of mapped reads which mapped to that variant) and Spearman rank-based correlation between replicates over only those variants that were detected in all four replicates (Table S19). While Pearson correlation between replicates and donors is over 90% for most pairwise comparisons, it is dominated by the magnitude of differences between the strongest- and weakest-detected variants. When the highest detected variants differ between samples, Pearson correlation plunges. Technical replicates are no better correlated than biological replicates, indicating substantial variability between library preps and sequencing reactions. This is not unexpected, given 93 total cycles of PCR amplification, as well as the known run-to-run variability in flow cell clustering efficiency when sequencing mixtures of inserts of different lengths. Spearman correlation generally tracks with Pearson correlation, but is less skewed by the magnitude of highest-detected variants. On the whole, Spearman correlation indicates a limited ability to accurately order variants according to detection level. We believe the highest resolution of detection level quantification possible from these sequencing experiments is the categorization of variants into several detection level ranges. We used ‘high’ (roughly 20% or more of mapped reads), ‘medium’ (roughly between 20% and 5%), ‘low’ (roughly between 5% and 1%), and ‘trace’ (roughly below 1%) categories. We then classified transcripts as consistent (in same detection level category) or inconsistent (different detection level categories) between donors. Allowing for observed variation between technical replicates, and considering ‘trace’ detection (typically fewer than 50, but always more than 5 reads) in some replicates consistent with no detection in others, the overwhelming majority of variants were detected at a consistent level between all three individuals (Table S9). Be believe the remaining, inconsistent variants reflect physiological differences between individuals.

### Plasmid construction

Full-length CaMKII variants were cloned into a pET vector containing N-terminal 6xHis followed by a SUMO tag. CaMKII chimeras were constructed as follows. The kinase/regulatory segment and hub domains of CaMKIIα and CaMKIIβ were amplified from full-length constructs. The kinase domain includes the regulatory segment of its corresponding kinase. Each domain was amplified with a 20 bps overhang that was the corresponding CaMKII gene. The chimeras were assembled using Gibson assembly.

### Protein expression and purification

All CaMKII variants used for this experiment were recombinantly expressed and purified according to (39). Plasmids encoding CaMKII variants were co-transformed with lambda phosphatase into Rosetta 2(DE3)pLysS competent cells (Millipore). Expression was induced overnight with 1mM Isopropyl β-D-1-thiogalactopyranoside (IPTG, GoldBio) and cultures were grown overnight at 18 °C. Cell pellets were resuspended in Buffer A (25 mM Tris-HCl pH 8.5, 150 mM KCl, 50 mM Imidazole, 10% glycerol) (Sigma) with 25 mM Magnesium chloride, containing a cocktail of protease inhibitors and DNase (AEBSF 0.2 mM, Leupeptin 0.005 mM, 1 µg/mL Pepstatin, 1 µg/mL Aprotinin, 0.1 mg/mL Trypsin Inhibitor, 0.5 mM Benzamidine, 1 µg/mL DNAse) (Sigma) and lysed. All subsequent purification steps were performed using an ÄKTA pure chromatography system at 4°C. Clarified lysate was loaded onto a 5 mL HisTrap FF NiNTA Sepharose column (GE), and eluted with a combination of 50% Buffer A and 50% Buffer B (25mM Tris-HCl pH 8.5, 150mM KCl, 1M imidazole, 10% glycerol) for a final concentration of 0.5 M imidazole. The protein was desalted of residual imidazole using a HiPrep 26/10 Desalting column, and His SUMO tags were cleaved with Ulp1 protease overnight at 4°C in Buffer C (25mM Tris-HCl pH 8.5, 150mM KCl, 2mM TCEP (GoldBio), 50mM Imidazole, 10% glycerol). Cleaved tags were separated by a subtractive NiNTA step. Next, an anion exchange step was done using a 5 mL HiTrap Q-FF and protein was eluted with a KCl gradient. Eluted proteins were concentrated and further purified in gel filtration buffer (25mM Tris-HCl pH 8.0, 150mM KCl, 1mM TCEP, 10% glycerol) using a Superose 6 Increase 10/300 size exclusion column. Pure fractions were then concentrated, aliquoted, flash frozen in liquid nitrogen and stored at −80°C until further use. All columns were purchased from GE.

Calmodulin (*Gallus gallus*) was recombinantly expressed from a pET-15b vector (generous gift from Angus Nairn) in BL21(DE3) cells (Millipore) and purified as previously described (40). To quantify the calmodulin concentration for making stocks to use in the kinase assays, we used circular dichroism on a Jasco J-1500 Spectrophotometer to make a measurement in triplicate for our purified sample scanning a wavelength spectrum between 250-215 nm to measure the characteristic wavelength of 222 nm according to (41).

Equation used to calculate calmodulin concentration:

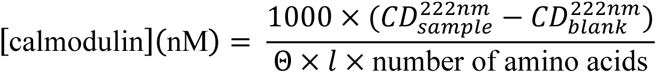

where circular dichroism at 222 nm (*CD*^*222nm*^) is expressed in mdeg, Θ is the molar ellipticity, and *l* is the path length in cm.

#### Coupled kinase assays

Kinase activity was monitored with previously described conditions (29, 42) using a Synergy H1 microplate reader (Biotek). The addition of calmodulin (concentrations ranging from 0nM-2μM) to the reaction was used to initiate CaMKII activity, after which absorbance was measured at 15 second intervals for 10 minutes. The change in absorbance over the course of the resulting time series of measurements was fit with a straight line (y = mx + c) to obtain a slope (m) proportional to the kinetic rate of the reaction. For each time series, slopes were fit to a sliding window of 5 points (1 minute 15 seconds) and the maximum observed slope was used to represent the kinetic rate of that reaction. Kinetic rates across a series of calmodulin concentrations were fit with the following equation:

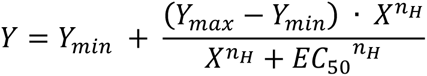

to obtain EC_50_ (defined as the calmodulin concentration needed to reach half the maximal reaction velocity) and cooperativity values (Hill coefficients, *n_H_*). 95% confidence intervals for fit parameters (EC_50_ and *n_H_*) were determined using the following bootstrap procedure. 10,000 replicate calmodulin concentration series were generated by randomly selecting one observed kinetic rate at each measured calmodulin concentration from the set of replicates for that variant. Each generated concentration series was fit with the equation above. Parameter values at the 2.5^th^ and 97.5^th^ quantiles of the 10,000 fits were taken as the boundaries of the 95% confidence interval.

| Assay Component | Final concentration in Master Mix |
| --- | --- |
| Calcium (CaCl<sub>2</sub>), Sigma | 0.2 mM |
| Pyruvate Kinase, Sigma | 55.7 U/ml |
| Lactate Dehydrogenase, Millipore Sigma | 78 U/ml |
| Adenosine triphosphate, Sigma | 0.5 mM |
| Phosphoenolpyruvate, Alfa Aesar | 1 mM |
| Nicotinamide adenine dinucleotide, Sigma | 0.2 mg/ml |
| Syntide, LifeTein | 0.3 mM |
| KCl, Sigma | 150 mM |
| Tris/MgCl<sub>2</sub> Buffer, Fisher Scientific | 50 mM/10 mM |

The final pH of each reaction was ∼7.5-8, and the final enzyme concentration was 133 nM.

#### Small Angle X-ray Scattering

Small Angle X-ray Scattering (SAXS) data were collected at the mail-in high-throughput SIBYLS beamline 12.3.1 at the Advanced Light Source (ALS) using a Pilatus3 2M pixel array detector (Dectris). 20 µl samples were loaded in a 1.5 mm thick chamber by an automated Tecan Freedom Evo liquid handling robot (43). Incident X-rays were tuned to a wavelength of 11keV/1.27Å, samples were exposed for 10 seconds and data collected every 0.1 or 0.33 sec. SAXS data were collected for CaMKIIα-0 at concentrations: 1, 3, 7.2 mg/mL, and CaMKIIβ-0 at 3, 7, 11.5 mg/mL. Frames were chosen using SAXS Frameslice (https://bl1231.als.lbl.gov/ran).

Data were then scaled and merged using the low q region from the lowest concentration datasets. R_g_ and D_max_ were determined using both Scatter 3.0 and primus (ATSAS 3.0).

#### Cryo-electron microscopy data collection and analysis

C-flat Holey Carbon grids with gold support (CF-1.2/1.3-3Au-50, EMS) were made hydrophilic by glow-discharging at 30 mA for 30 s using PELCO easiGlow (Ted Pella). 3 µl of purified CaMKII at 0.5-5 mg/ml was applied to these grids at ∼96% relative humidity, blotted for 5 s and plunge-frozen into liquid ethane using Cryoplunge 3 (Gatan).

Frozen hydrated samples were imaged on a FEI Talos Arctica at 200 kV with K2 Summit direct detection camera in super-resolution mode with a total exposure dose of around 52 electrons. Thirty-five frames per movie were collected at a magnification corresponding to 0.455 Å per pixel using SerialEM (44). Micrographs (6510) were collected at defocus values ranging from −0.5 to −2.5 µm. Movie frames were motion-corrected and dose-weighted by MotionCor2 (45), sampled to 0.91 Å per pixel and contrast transfer function (CTF) parameters were estimated by CTFFIND4 (46). Particle picking was carried out using crYOLO giving 1,500,734 initial particles. Initial *ab-initio* models were generated in Relion (47). Following successive rounds of 2D and 3D classification 290,980 particles were selected. An additional rounds of 3D classification led to the reconstructions of 4.8 Å from 160,211 particles (reconstruction Class A), of 6.6 Å from 40,329 particles (reconstruction Class B, fig. S4).

Maps used for figures were filtered according to local resolution with b-factor sharpening within Relion. Models were fit into the map using Chimera (48) and Coot (49) and rigid body refined with PHENIX (50).

## Acknowledgments

We thank Florian Heyd and Andreas Franz for helpful discussions on data and the manuscript. We also thank Peter Chien, Eric Strieter, Scott Garman for helpful discussions. We thank Greg Hura and Kathryn Burnett for SAXS data collection and help with analysis. SAXS data was collected at the Advanced Light Source (ALS), SIBYLS beamline on behalf of US DOE-BER, through the Integrated Diffraction Analysis Technologies (IDAT) program. Additional support comes from the NIGMS project ALS-ENABLE (P30 GM124169) and a High-End Instrumentation Grant S10OD018483. Cryo-EM data were collected at the Harvard Cryo-EM Center for Structural Biology. We are grateful for support from Sarah Sterling, Richard Walsh, Shaun Rawson and Zongli Li for cryo-EM data collection.

## Author contributions

RS and ND designed experiments, collected and analyzed data, wrote and edited manuscript. MD, RB, AT, BP, JW, SB, LC collected data. MS and LC designed experiments, analyzed data, wrote and edited manuscript.

## Data and Materials Availability

Cryo-EM maps and models to be deposited into EMDB and the PDB.

## Supplementary Materials

**Fig. S1.**
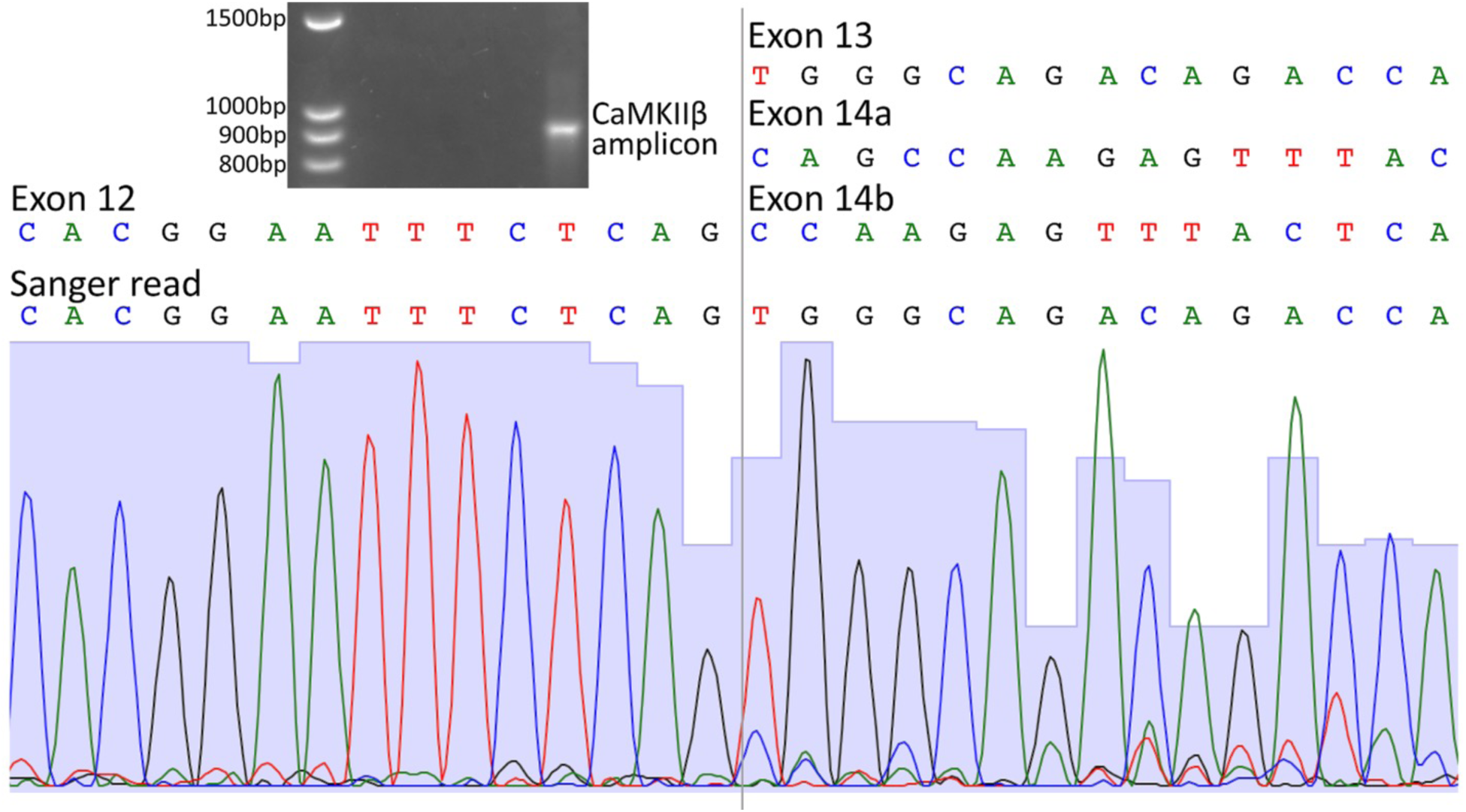
PCR amplicons from multiple CaMKIIβ variants do not separate on agarose gel. A PCR reaction with CaMKIIβ-specific primers was loaded onto a 2.5% agarose gel and run at 10V for roughly 24hrs (pictured below) for optimal band separation. The single visible band was excised, purified, and Sanger-sequenced. A fragment of the resulting sequencing trace is shown, with height of grey background representing Phred quality scores. The first 15bp of the fragment (left of grey line) map to the last 15bp of CaMKIIβ constitutive exon 12, incorporated into every CaMKIIβ transcript. The last 15bp of the fragment (right of grey line) show a dramatic increase of secondary peak heights, with a corresponding drop-off in Phred scores. The sequence defined by primary peaks (TGGGCAGACAGACCA) maps to CaMKIIβ linker exon 13. Variants β13,14b,16,17,18 (44.8%), β13,14b,16,18 (5.9%), β13,14b,17,18 (4.0%), β13,14b,16,17,18-short-hub (3.7%) accounted for roughly 58.4% of mapped reads from the CaMKIIβ Illumina library. Variants in which exon 14a (CAGCCAAGAGTTTAC) followed exon 12, such as β14a,16,18 (13.8%) and β14a,16,17,18 (9.1%), accounted for roughly 30% of mapped reads. Variants in which exon 14b (CCAAGAGTTTACTCA) followed exon 12, such as β14a, 16,17,18 (7.1%), β14a,16,18 (6.3%), and β14a,17,18 (1.2%) accounted for roughly 14% of mapped reads. Secondary peaks corresponding to exons 14a and 14b are clearly visible in the last 15bp of the Sanger sequencing fragment.

**Fig. S2.**
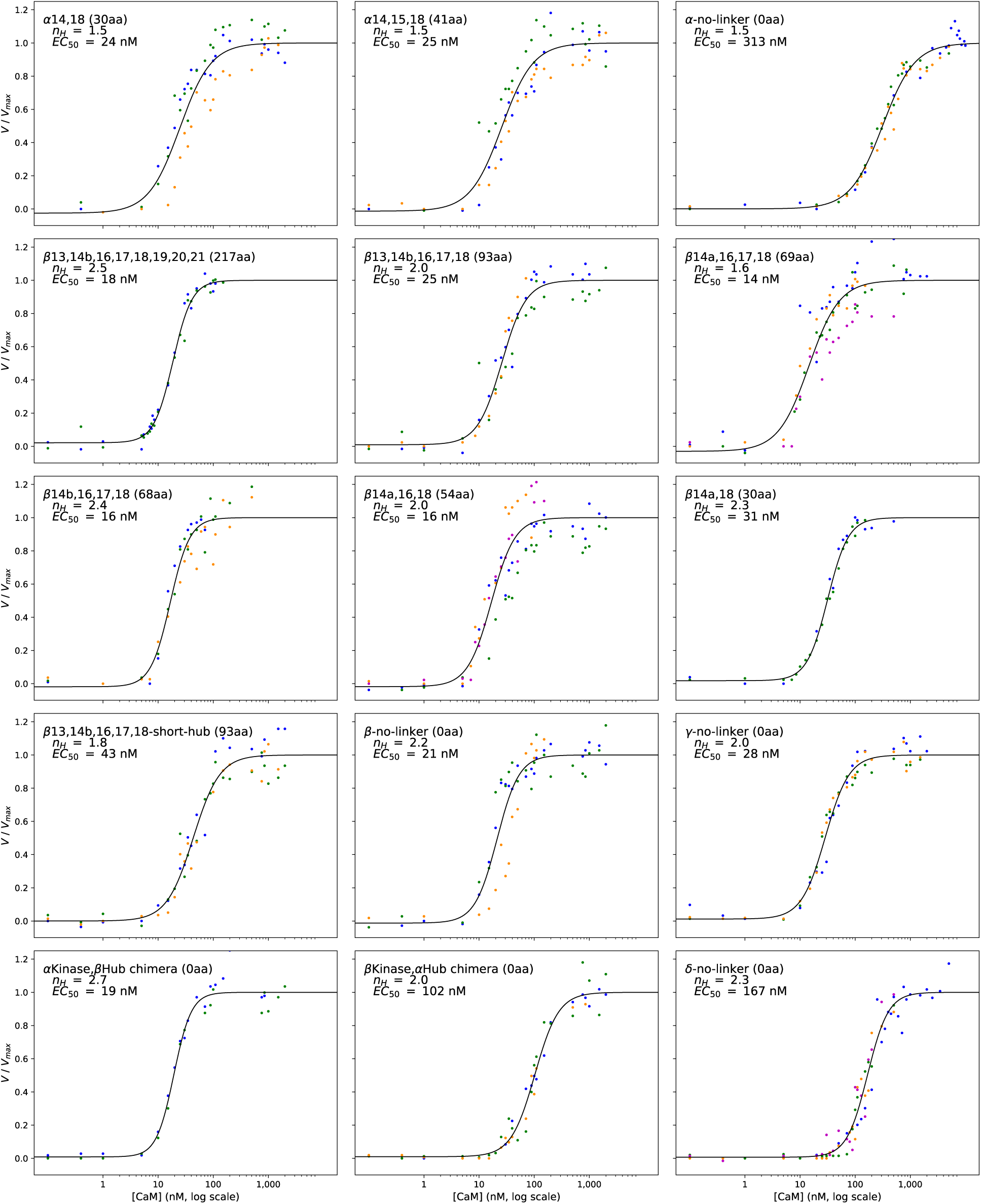
Data and EC_50_ fits for all tested CaMKII variants. Data point color reflects replicates.

**Fig. S3.**
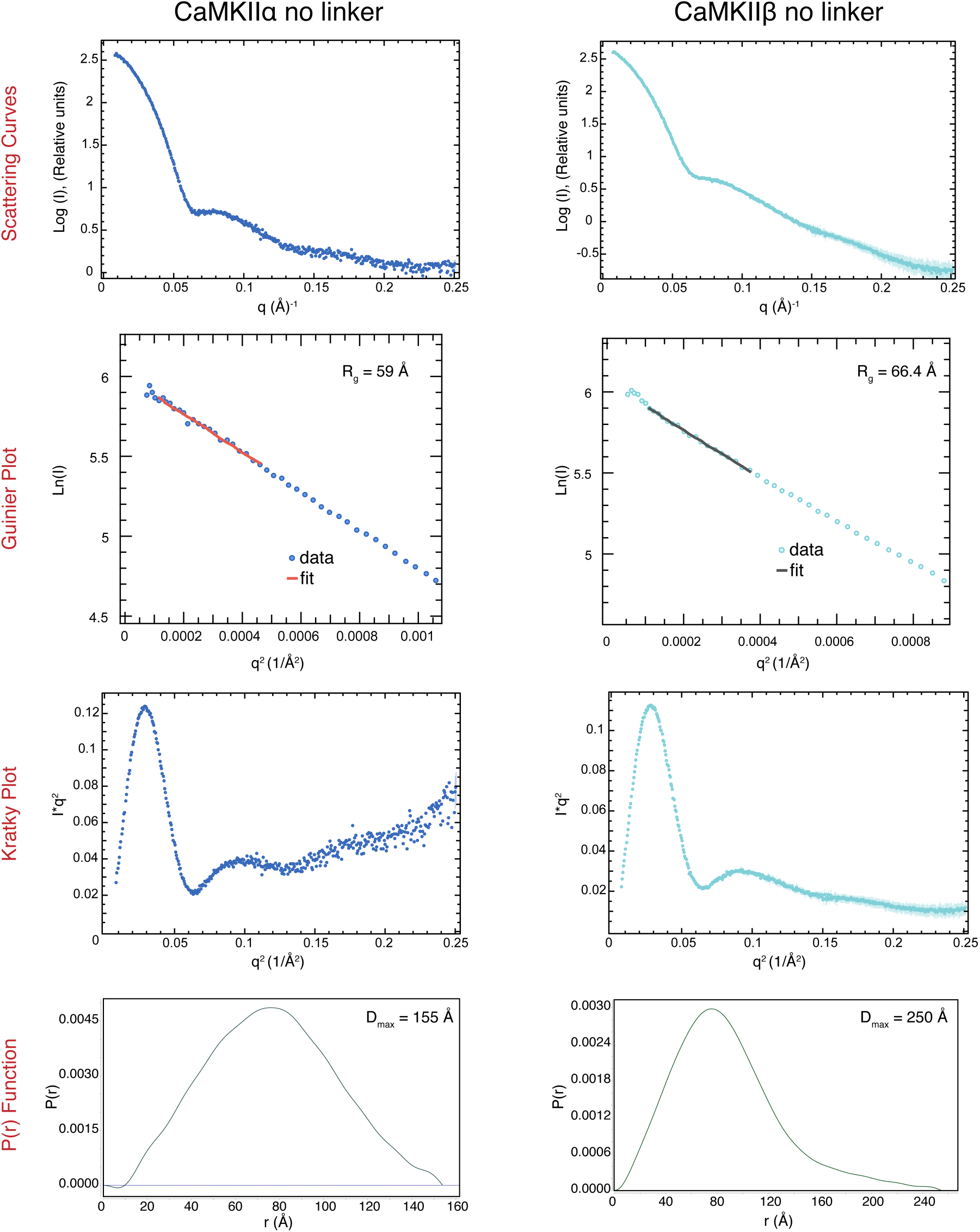
SAXS data and analyses of CaMKIIα-0 and CaMKIIβ-0. Scattering curves, Guinier plots and Kratky plots were made using Primus (ATSAS 3.0). Pair distribution functions were made using Scatter 3.0.

**Fig. S4.**
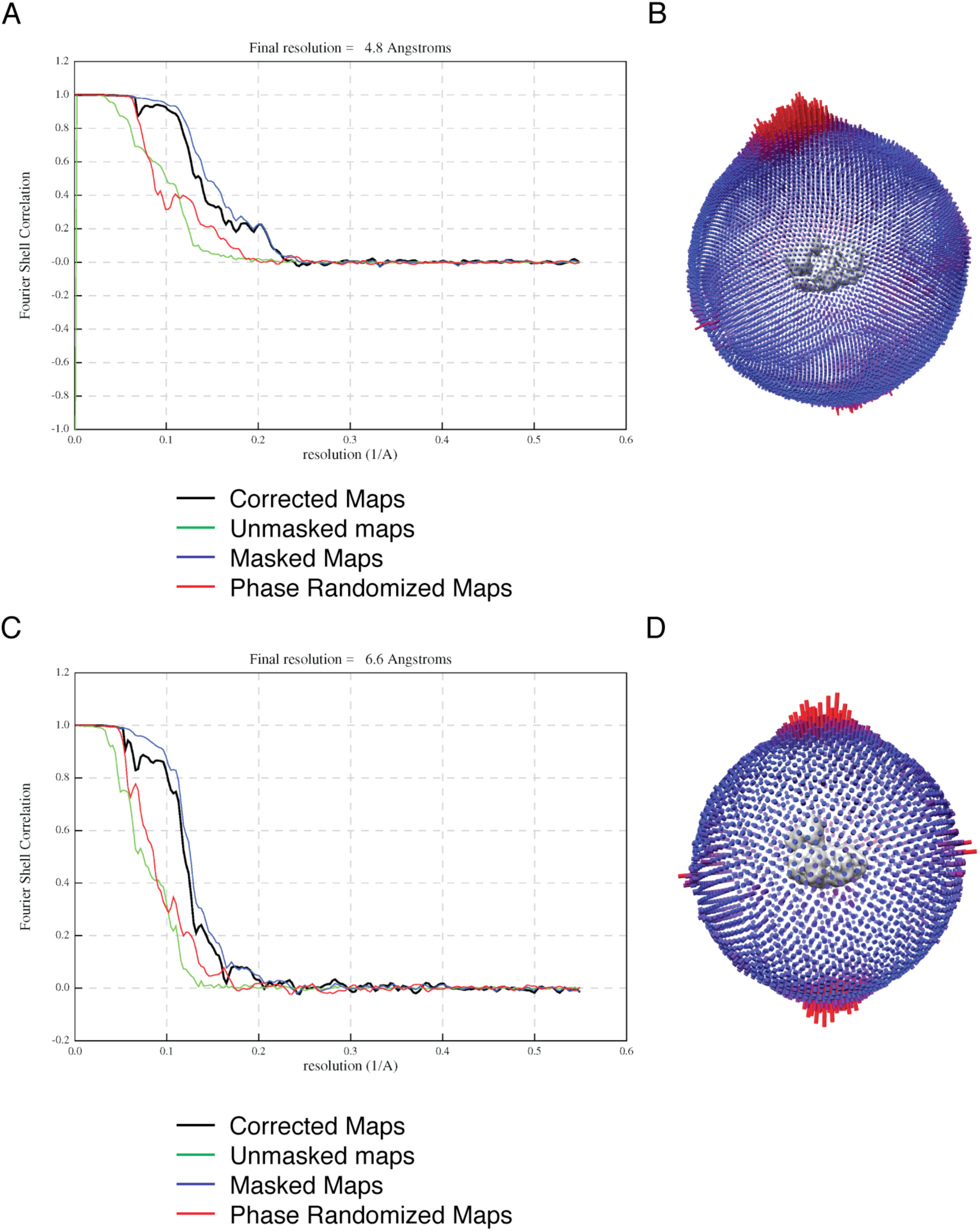
Cryo-EM data. (A) Fourier shell correlation (FSC) curves obtained from Relion 3 postprocess job for Class A reconstruction. The resolution is estimated at FSC = 0.143. (B) Angular distribution of the final reconstruction obtained from Relion for Class A. (C) Fourier shell correlation (FSC) curves obtained from Relion 3 postprocess job for Class B reconstruction. The resolution is estimated at FSC = 0.143. (D) (B) Angular distribution of the final reconstruction obtained from Relion for Class B.

**Table S1.**
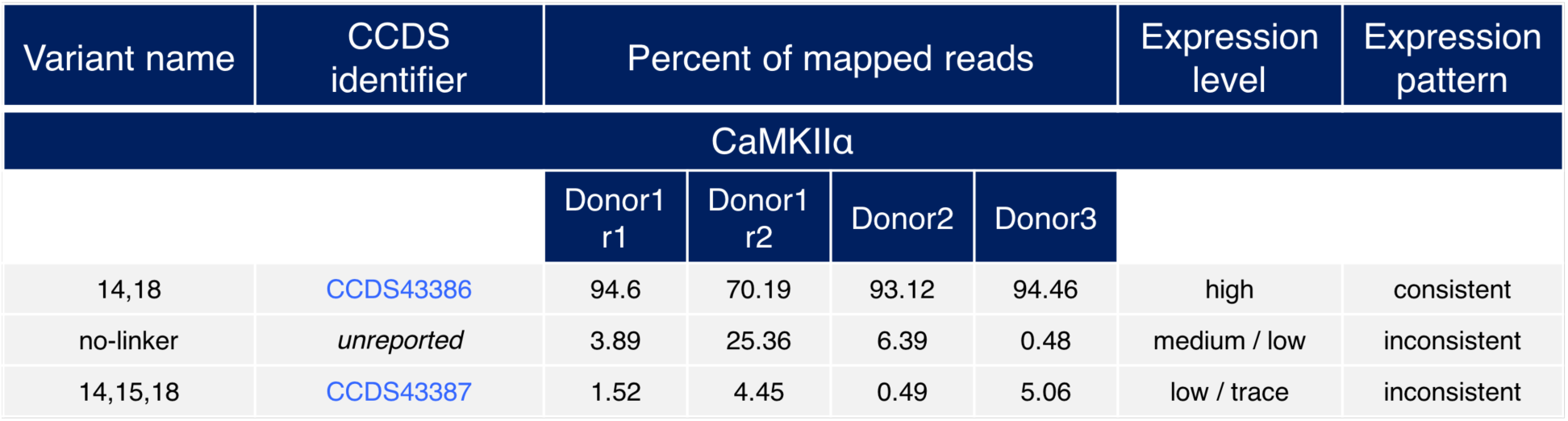
Detected CaMKIIα splice variants

| Variant name | CCDS identifier | Percent of mapped reads |  |  |  | Expression level | Expression pattern |
| --- | --- | --- | --- | --- | --- | --- | --- |
| CaMKIIα | | | | | | | |
|  |  | Donor1<br>r1 | Donor1<br>r2 | Donor2 | Donor3 |  |  |
| 14,18 | CCDS43386 | 94.6 | 70.19 | 93.12 | 94.46 | high | consistent |
| no-linker | *unreported* | 3.89 | 25.36 | 6.39 | 0.48 | medium / low | inconsistent |
| 14,15,18 | CCDS43387 | 1.52 | 4.45 | 0.49 | 5.06 | low / trace | inconsistent |

**Table S2.** Read counts of CaMKIIα splice variants from Donor 1

| CaMKIIα | | | | |
| --- | --- | --- | --- | --- |
| Donor 1 |  |  |  |  |
| Replicate 1 |  |  | Replicate 2 |  |
| Variant | Read count | % mapped | Read count | % mapped |
| 14,18 | 183118 | 94.6 | 81521 | 70.2 |
| no-linker | 7527 | 3.89 | 29449 | 25.4 |
| 14,15,18 | 2941 | 1.52 | 5170 | 4.45 |
| **Total** | **193586** |  | **116140** |  |

**Table S3.** Read counts of CaMKIIα splice variants from Donor 2

| CaMKIIα | | |
| --- | --- | --- |
| Donor 2 |  |  |
| Variant | Read count | % mapped |
| 14,18 | 81002 | 93.1 |
| no-linker | 5558 | 6.39 |
| 14,15,18 | 426 | 0.49 |
| **Total** | **86986** |  |

**Table S4.** . Read counts of CaMKIIα splice variants from Donor 3

| CaMKIIα | | |
| --- | --- | --- |
| Donor 3 |  |  |
| Variant | Read count | % mapped |
| 14,18 | 53383 | 94.5 |
| no-linker | 2862 | 5.06 |
| 14,15,18 | 271 | 0.48 |
| **Total** | **56516** |  |

**Table S5.** Detected CaMKIIβ splice variants

| Variant name | CCDS identifier | Percent of mapped reads |  |  |  | Expression level | Expression pattern |
| --- | --- | --- | --- | --- | --- | --- | --- |
| CaMKIIβ |  |  |  |  |  |  |  |
|  |  | Donor1<br>r1 | Donor1<br>r2 | Donor2 | Donor3 |  |  |
| 13,14b,16,17,18,full | CCDS5484 | 44.83 | 56.31 | 54.02 | 36.63 | high | consistent |
| 14a,16,18,full | CCDS5486 | 13.78 | 5.85 | 6.59 | 7.40 | medium | consistent |
| 14a,16,17,18,full | CCDS5485 | 9.11 | 6.99 | 6.06 | 9.34 | medium | consistent |
| 14b,16,17,18,full | CCDS43573 | 7.08 | 5.73 | 3.95 | 6.50 | medium | consistent |
| 14b,17,18,full | *unreported* | 1.16 | 5.74 | 14.35 | 4.46 | medium / low | inconsistent |
| 13,14b,16,18,full | *unreported* | 5.89 | 5.13 | 2.62 | 6.00 | medium | consistent |
| 13,14b,17,18,full | *unreported* | 3.97 | 3.51 | 1.92 | 10.30 | medium / low | inconsistent |
| 14b,16,18,full | *unreported* | 6.28 | 2.08 | 1.50 | 3.21 | low | consistent |
| 13,14b,18,full | *unreported* | 0.63 | 4.29 | 4.25 | 1.28 | low | consistent |
| 14a,17,18,full | *unreported* | 0.83 | 1.35 | 2.39 | 2.32 | low | consistent |
| 13,14b,16,17,18,short | *unreported* | 3.72 | 1.12 | 0.03 | 2.56 | low / trace | inconsistent |
| 14a,18,full | CCDS5488 | 0.51 | 0.58 | 1.95 | 1.83 | low | consistent |
| 14b,18,full | *unreported* | 0.24 | 0.26 | 0.34 | 0.53 | trace | consistent |
| 13,14a,16,17,18,full | *unreported* | 0.28 | 0.23 | 0.02 | 0.10 | trace | consistent |
| 14b,16,17,18,short | *unreported* | 0.43 | 0.13 | N | 0.43 | trace | consistent |
| 13,14b,16,18,short | *unreported* | 0.27 | 0.10 | N | 0.45 | trace | consistent |
| 14a,16,17,18,short | *unreported* | 0.21 | 0.08 | N | 0.66 | trace | consistent |
| 13,14b,17,18,short | CCDS5487 | 0.10 | 0.08 | N | 0.83 | trace | consistent |
| 14a,16,18,short | *unreported* | 0.21 | 0.08 | N | 0.35 | trace | consistent |
| 14b,16,18,short | *unreported* | 0.07 | 0.13 | N | 0.23 | trace | consistent |
| no-linker,full | CCDS5489 | 0.13 | N | N | 3.67 | low / trace | inconsistent |
| 14a,17,18,short | *unreported* | 0.11 | N | N | 0.15 | trace | consistent |
| 13,full | *unreported* | 0.04 | N | N | 0.08 | trace | consistent |
| 14b,17,18,short | *unreported* | N | 0.07 | N | 0.22 | trace | consistent |
| 13,14a,16,18,full | *unreported* | 0.12 | N | N | N | trace | consistent |
| 16,17,18,full | *unreported* | N | 0.16 | N | N | trace | consistent |
| 13,14b,18,short | *unreported* | N | 0.04 | N | N | trace | consistent |
| 18,full | *unreported* | N | N | N | 0.22 | trace | consistent |
| no-linker,short | *unreported* | N | N | N | 0.15 | trace | consistent |
| 14b,18,short | *unreported* | N | N | N | 0.08 | trace | consistent |

**Table S6.** Read counts of CaMKIIβ splice variants from Donor 1

| CaMKIIβ | | | | | |
| --- | --- | --- | --- | --- | --- |
| Donor 1 |  |  |  |  |  |
| Replicate 1 |  |  | Replicate 2 |  |  |
| Variant | Read count | % mapped | Variant | Read count | % mapped |
| 13,14b,16,17,18,full | 65577 | 44.83 | 13,14b,16,17,18,full | 23166 | 56.3 |
| 14a,16,18,full | 20153 | 13.8 | 14a,16,17,18,full | 2875 | 6.99 |
| 14a,16,17,18,full | 13327 | 9.11 | 14a,16,18,full | 2405 | 5.85 |
| 14b,16,17,18,full | 10362 | 7.08 | 14b,17,18,full | 2363 | 5.74 |
| 14b,16,18,full | 9192 | 6.28 | 14b,16,17,18,full | 2358 | 5.73 |
| 13,14b,16,18,full | 8616 | 5.89 | 13,14b,16,18,full | 2109 | 5.13 |
| 13,14b,17,18,full | 5805 | 3.97 | 13,14b,18,full | 1765 | 4.29 |
| 13,14b,16,17,18,short | 5442 | 3.72 | 13,14b,17,18,full | 1446 | 3.51 |
| 14b,17,18,full | 1700 | 1.16 | 14b,16,18,full | 855 | 2.08 |
| 14a,17,18,full | 1218 | 0.83 | 14a,17,18,full | 555 | 1.35 |
| 13,14b,18,full | 928 | 0.63 | 13,14b,16,17,18,short | 459 | 1.12 |
| 14a,18,full | 750 | 0.51 | 14a,18,full | 237 | 0.58 |
| 14b,16,17,18,short | 634 | 0.43 | 14b,18,full | 105 | 0.26 |
| 13,14a,16,17,18,full | 412 | 0.28 | 13,14a,16,17,18,full | 94 | 0.23 |
| 13,14b,16,18,short | 400 | 0.27 | 16,17,18,full | 64 | 0.16 |
| 14b,18,full | 352 | 0.24 | 14b,16,18,short | 52 | 0.13 |
| 14a,16,17,18,short | 309 | 0.21 | 14b,16,17,18,short | 52 | 0.13 |
| 14a,16,18,short | 303 | 0.21 | 13,14b,16,18,short | 41 | 0.10 |
| no-linker,full | 185 | 0.13 | 13,14b,17,18,short | 33 | 0.08 |
| 13,14a,16,18,full | 169 | 0.12 | 14a,16,18,short | 33 | 0.08 |
| 14a,17,18,short | 155 | 0.11 | 14a,16,17,18,short | 32 | 0.08 |
| 13,14b,17,18,short | 148 | 0.10 | 14b,17,18,short | 27 | 0.07 |
| 14b,16,18,short | 104 | 0.07 | 13,14b,18,short | 17 | 0.04 |
| 13,full | 52 | N | **Total** | **41143** |  |
| **Total** | **146293** |  |  |  |  |

**Table S7.** Read counts of CaMKIIβ splice variants from Donor 2

| CaMKIIβ | | |
| --- | --- | --- |
| Donor 2 |  |  |
| Variant | Read count | % mapped |
| 13,14b,16,17,18,full | 28645 | 54.0 |
| 14b,17,18,full | 7609 | 14.4 |
| 14a,16,18,full | 3495 | 6.59 |
| 14a,16,17,18,full | 3215 | 6.06 |
| 13,14b,18,full | 2255 | 4.25 |
| 14b,16,17,18,full | 2095 | 3.95 |
| 13,14b,16,18,full | 1390 | 2.62 |
| 14a,17,18,full | 1269 | 2.39 |
| 14a,18,full | 1036 | 1.95 |
| 13,14b,17,18,full | 1016 | 1.92 |
| 14b,16,18,full | 795 | 1.50 |
| 14b,18,full | 178 | 0.34 |
| 13,14b,16,17,18,short | 15 | 0.03 |
| 13,14a,16,17,18,full | 11 | 0.02 |
| **Total** | **53024** |  |

**Table S8.** Read counts of CaMKIIβ splice variants from Donor 3

| CaMKIIβ | | |
| --- | --- | --- |
| Donor 3 |  |  |
| Variant | Read count | % mapped |
| 13,14b,16,17,18,full | 18165 | 36.6 |
| 13,14b,17,18,full | 5107 | 10.3 |
| 14a,16,17,18,full | 4629 | 9.34 |
| 14a,16,18,full | 3669 | 7.40 |
| 14b,16,17,18,full | 3225 | 6.50 |
| 13,14b,16,18,full | 2977 | 6.00 |
| 14b,17,18,full | 2212 | 4.46 |
| no-linker,full | 1821 | 3.67 |
| 14b,16,18,full | 1591 | 3.21 |
| 13,14b,16,17,18,short | 1270 | 2.56 |
| 14a,17,18,full | 1149 | 2.32 |
| 14a,18,full | 907 | 1.83 |
| 13,14b,18,full | 637 | 1.28 |
| 13,14b,17,18,short | 413 | 0.83 |
| 14a,16,17,18,short | 328 | 0.66 |
| 14b,18,full | 263 | 0.53 |
| 13,14b,16,18,short | 224 | 0.45 |
| 14b,16,17,18,short | 215 | 0.43 |
| 14a,16,18,short | 172 | 0.35 |
| 14b,16,18,short | 116 | 0.23 |
| 18,full | 110 | 0.22 |
| 14b,17,18,short | 108 | 0.22 |
| 14a,17,18,short | 75 | 0.15 |
| no-linker,short | 72 | 0.15 |
| 13,14a,16,17,18,full | 49 | 0.10 |
| 13,full | 42 | 0.08 |
| 14b,18,short | 40 | 0.08 |
| **Total** | **49586** |  |

**Table S9.**
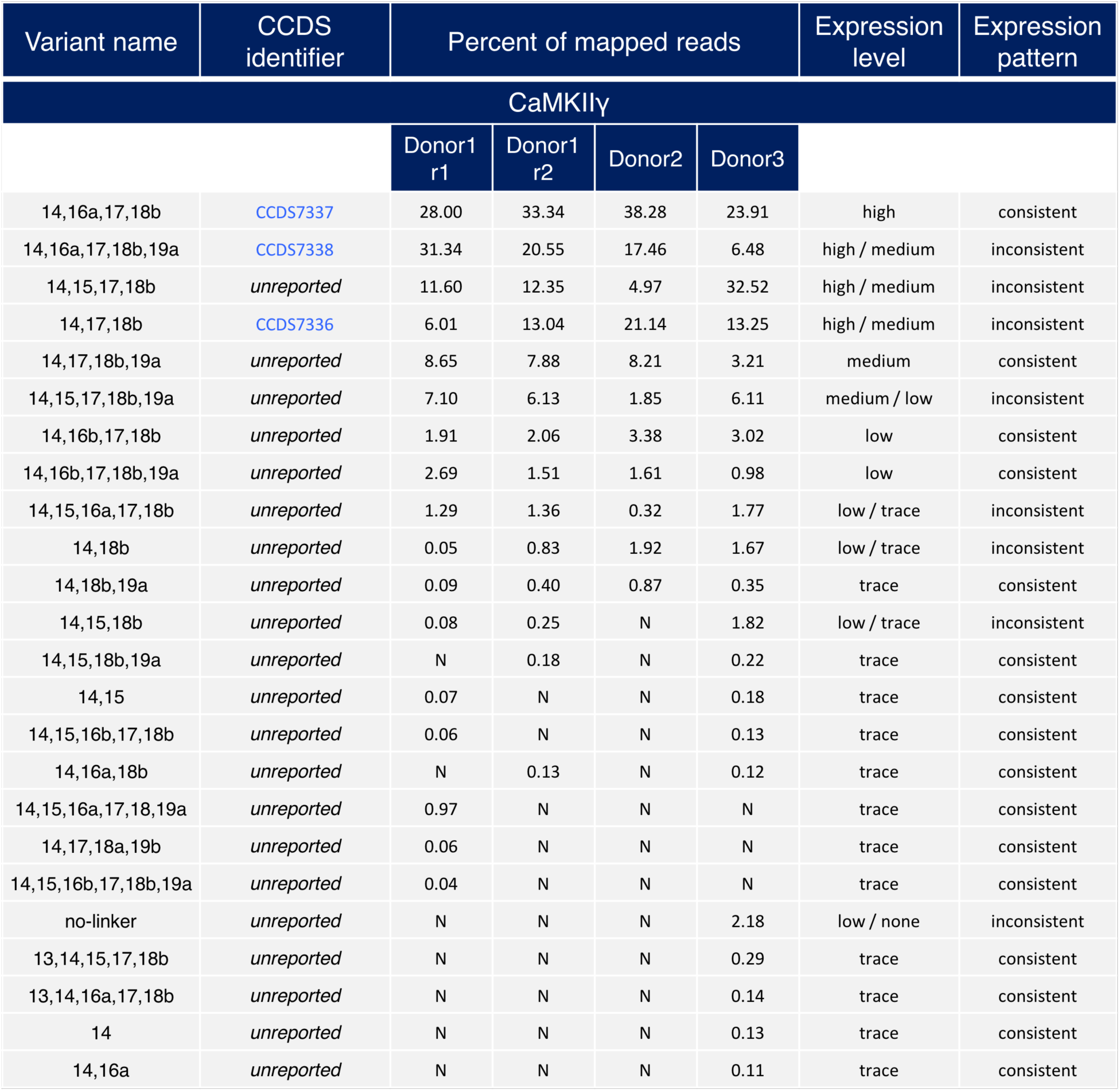
Detected CaMKIIγ splice variants

**Table S10.** Read counts of CaMKIIγ splice variants from Donor 1

| CaMKIIγ | | | | | |
| --- | --- | --- | --- | --- | --- |
| Donor 1 |  |  |  |  |  |
| Replicate 1 |  |  | Replicate 2 |  |  |
| Variant | Read count | % mapped | Variant | Read count | % mapped |
| 14,16a,17,18b,19a | 53510 | 31.3 | 14,16a,17,18b | 11440 | 33.3 |
| 14,16a,17,18b | 47800 | 28.0 | 14,16a,17,18b,19a | 7051 | 20.5 |
| 14,15,17,18b | 19801 | 11.6 | 14,17,18b | 4475 | 13.0 |
| 14,17,18b,19a | 14773 | 8.65 | 14,15,17,18b | 4237 | 12.3 |
| 14,15,17,18b,19a | 12127 | 7.10 | 14,17,18b,19a | 2705 | 7.88 |
| 14,17,18b | 10268 | 6.01 | 14,15,17,18b,19a | 2102 | 6.13 |
| 14,16b,17,18b,19a | 4586 | 2.69 | 14,16b,17,18b | 708 | 2.06 |
| 14,16b,17,18b | 3253 | 1.91 | 14,16b,17,18b,19a | 517 | 1.51 |
| 14,15,16a,17,18b | 2196 | 1.29 | 14,15,16a,17,18b | 465 | 1.36 |
| 14,15,16a,17,18b,19a | 1662 | 0.97 | 14,18b | 285 | 0.83 |
| 14,18b,19a | 160 | 0.09 | 14,18b,19a | 136 | 0.40 |
| 14,15,18b | 133 | 0.08 | 14,15,18b | 87 | 0.25 |
| 14,15 | 111 | 0.07 | 14,15,18b,19a | 61 | 0.18 |
| 14,15,16b,17,18b | 100 | 0.06 | 14,16a,18b | 45 | 0.13 |
| 14,17,18a,19b | 96 | 0.06 | **Total** | **34314** |  |
| 14,18b | 81 | 0.05 |  |  |  |
| 14,15,16b,17,18b,19a | 72 | 0.04 |  |  |  |
| **Total** | **170729** |  |  |  |  |

**Table S11.** Read counts of CaMKIIγ splice variants from Donor 2

| CaMKIIγ | | |
| --- | --- | --- |
| Donor 2 |  |  |
| Variant | Read count | % mapped |
| 14,16a,17,18b | 11647 | 38.3 |
| 14,17,18b | 6433 | 21.1 |
| 14,16a,17,18b,19a | 5314 | 17.5 |
| 14,17,18b,19a | 2498 | 8.21 |
| 14,15,17,18b | 1512 | 4.97 |
| 14,16b,17,18b | 1029 | 3.38 |
| 14,18b | 584 | 1.92 |
| 14,15,17,18b,19a | 562 | 1.85 |
| 14,16b,17,18b,19a | 489 | 1.61 |
| 14,18b,19a | 265 | 0.87 |
| 14,15,16a,17,18b | 96 | 0.32 |
| **Total** | **30429** |  |

**Table S12.** Read counts of CaMKIIγ splice variants from Donor 3

| CaMKIIγ | | |
| --- | --- | --- |
| Donor 3 |  |  |
| Variant | Read count | % mapped |
| 14,15,17,18b | 9131 | 32.5 |
| 14,16a,17,18b | 6714 | 23.9 |
| 14,17,18b | 3719 | 13.2 |
| 14,15,17,18b,19a | 1716 | 6.11 |
| 14,16a,17,18b,19a | 1820 | 6.48 |
| 14,17,18b,19a | 900 | 3.21 |
| 14,16b,17,18b | 849 | 3.02 |
| no-linker | 611 | 2.18 |
| 14,15,18b | 512 | 1.82 |
| 14,15,16a,17,18b | 497 | 1.77 |
| 14,18b | 469 | 1.67 |
| 14,15,17,18b | 395 | 1.41 |
| 14,16b,17,18b,19a | 274 | 0.98 |
| 14,18b,19a | 99 | 0.35 |
| 13,14,15,17,18b | 82 | 0.29 |
| 14,15,18b,19a | 63 | 0.22 |
| 14,15 | 51 | 0.18 |
| 13,14,16a,17,18b | 38 | 0.14 |
| 14,15,16b,17,18b | 37 | 0.13 |
| 14 | 36 | 0.13 |
| 14,16a,18b | 33 | 0.12 |
| 14,16a | 30 | 0.11 |
| **Total** | **30429** |  |

**Table S13.** Detected CaMKIIδ splice variants

| Variant name | CCDS identifier | Percent of mapped reads |  |  |  | Expression level | Expression pattern |
| --- | --- | --- | --- | --- | --- | --- | --- |
| CaMKIIδ |  |  |  |  |  |  |  |
|  |  | Donor1 r1 | Donor1 r2 | Donor2 | Donor3 |  |  |
| 6v1,14b,18 | CCDS82950 | 53.13 | 44.70 | 40.15 | 26.38 | high | consistent |
| 6v1,14a,18 |  | 29.04 | 33.59 | 21.63 | 33.65 | high | consistent |
| 6v1,14a,16,17,18 | CCDS3703<br>CCDS3704<br>CCDS43263 | 7.62 | 11.56 | 20.78 | 15.59 | medium | consistent |
| 6v2,14b,18 | *unreported* | 4.70 | 0.89 | 5.74 | 2.86 | low | consistent |
| 6v2,14a,18 | *unreported* | 1.44 | 1.04 | 1.72 | 1.19 | low | consistent |
| 6v1,14b,16,17,18 | *unreported* | 0.10 | 0.61 | 0.07 | 1.09 | trace | consistent |
| 6v1,14a,17,18 | *unreported* | 1.32 | 3.45 | N | 9.53 | medium / low / none | inconsistent |
| 6v1,14b,17,18 | CCDS47127 | 0.96 | 1.42 | N | 1.27 | low / none | inconsistent |
| 6v1,14a,15,17,18 | *unreported* | 0.11 | N | 6.18 | 2.35 | low / trace | inconsistent |
| 6v2,14a,17,18 | *unreported* | 0.66 | 0.07 | N | 1.26 | low / none | inconsistent |
| 6v1,14,15,16,17,18 | *unreported* | 0.03 | 0.20 | N | 0.17 | trace | consistent |
| 6v2,14b,16,17,18 | *unreported* | N | 0.03 | 1.92 | 0.07 | low / trace | inconsistent |
| 6v2,14a,16,17,18 | *unreported* | N | N | 2.15 | 3.26 | low / none | inconsistent |
| 6v1,14a | *unreported* | 0.38 | 1.23 | N | N | trace | consistent |
| 6v1,14b | *unreported* | 0.51 | 0.75 | N | N | trace | consistent |
| 6v2,14a,15,17,18 | *unreported* | N | N | 0.73 | 0.40 | trace | consistent |
| 6v1,14b,15,17,18 | *unreported* | N | N | 0.65 | 0.37 | trace | consistent |
| 6v1,14a,15,18 | *unreported* | N | 0.30 | N | 0.30 | trace | consistent |
| 6v1,14b,15,18 | *unreported* | N | 0.08 | N | N | trace | consistent |
| 6v2,14b,17,18 | *unreported* | N | 0.04 | N | N | trace | consistent |
| 6v1,14a,16,18 | *unreported* | N | N | N | 0.17 | trace | consistent |
| 6v2,14b,15,17,18 | *unreported* | N | N | N | 0.07 | trace | consistent |

**Table S14.** Read counts of CaMKIIδ splice variants from Donor 1

| CaMKIIδ |  |  |  |  |  |
| --- | --- | --- | --- | --- | --- |
| Donor 1 |  |  |  |  |  |
| Replicate 1 |  |  | Replicate 2 |  |  |
| Variant | Read count | % mapped | Variant | Read count | % mapped |
| 6v1,14b,18 | 45214 | 53.1 | 6v1,14b,18 | 12388 | 44.7 |
| 6v1,14a,18 | 24712 | 29.0 | 6v1,14a,18 | 9310 | 33.6 |
| 6v1,14a,16,17,18 | 6483 | 7.62 | 6v1,14a,16,17,18 | 3204 | 11.6 |
| 6v2,14b,18 | 4001 | 4.70 | 6v1,14a,17,18 | 957 | 3.45 |
| 6v2,14a,18 | 1225 | 1.44 | 6v1,14b,17,18 | 393 | 1.42 |
| 6v1,14a,17,18 | 1121 | 1.32 | 6v1,14a | 341 | 1.23 |
| 6v1,14b,17,18 | 820 | 0.96 | 6v2,14a,18 | 289 | 1.04 |
| 6v2,14a,17,18 | 563 | 0.66 | 6v2,14b,18 | 247 | 0.89 |
| 6v1,14b | 431 | 0.51 | 6v1,14b | 209 | 0.75 |
| 6v1,14a | 326 | 0.38 | 6v1,14b,16,17,18 | 170 | 0.61 |
| 6v1,14a,15,17,18 | 90 | 0.11 | 6v1,14a,15,18 | 84 | 0.30 |
| 6v1,14b,16,17,18 | 84 | 0.10 | 6v1,14b,15,18 | 21 | 0.08 |
| **Total** | **85070** |  | 6v2,14a,17,18 | 20 | 0.07 |
|  |  |  | 6v2,14b,17,18 | 10 | 0.04 |
|  |  |  | 6v2,14b,16,17,18 | 9 | 0.03 |
|  |  |  | **Total** | **27660** |  |

**Table S15.** Read counts of CaMKIIδ splice variants from Donor 2

| CaMKIIδ |  |  |
| --- | --- | --- |
| Donor 2 |  |  |
| Variant | Read count | % mapped |
| 6v1,14b,18 | 7318 | 40.1 |
| 6v1,14a,18 | 3942 | 21.6 |
| 6v1,14a,16,17,18 | 3788 | 20.8 |
| 6v1,14a,15,17,18 | 1126 | 6.18 |
| 6v2,14b,18 | 1047 | 5.74 |
| 6v2,14a,16,17,18 | 392 | 2.15 |
| 6v2,14a,18 | 314 | 1.72 |
| 6v2,14a,15,17,18 | 133 | 0.73 |
| 6v1,14b,15,17,18 | 118 | 0.65 |
| 6v2,14b,16,17,18 | 36 | 0.20 |
| 6v1,14b,16,17,18 | 13 | 0.07 |
| **Total** | **18227** |  |

**Table S16.** Read counts of CaMKIIδ splice variants from Donor 3

| CaMKIIδ | | |
| --- | --- | --- |
| Donor 3 |  |  |
| Variant | Read count | % mapped |
| 6v1,14a,18 | 5094 | 33.6 |
| 6v1,14b,18 | 3993 | 26.4 |
| 6v1,14a,16,17,18 | 2360 | 15.6 |
| 6v1,14a,17,18 | 1443 | 9.53 |
| 6v2,14a,16,17,18 | 493 | 3.26 |
| 6v2,14b,18 | 433 | 2.86 |
| 6v1,14a,15,17,18 | 356 | 2.35 |
| 6v1,14b,17,18 | 192 | 1.27 |
| 6v2,14a,17,18 | 190 | 1.26 |
| 6v2,14a,18 | 180 | 1.19 |
| 6v1,14b,16,17,18 | 165 | 1.09 |
| 6v2,14a,15,17,18 | 61 | 0.40 |
| 6v1,14b,15,17,18 | 56 | 0.37 |
| 6v1,14a,15,18 | 45 | 0.30 |
| 6v1,14a,16,18 | 25 | 0.17 |
| 6v2,14b,16,17,18 | 11 | 0.07 |
| 6v2,14b,15,17,18 | 11 | 0.07 |
| **Total** | **15108** |  |

**Table S17.**
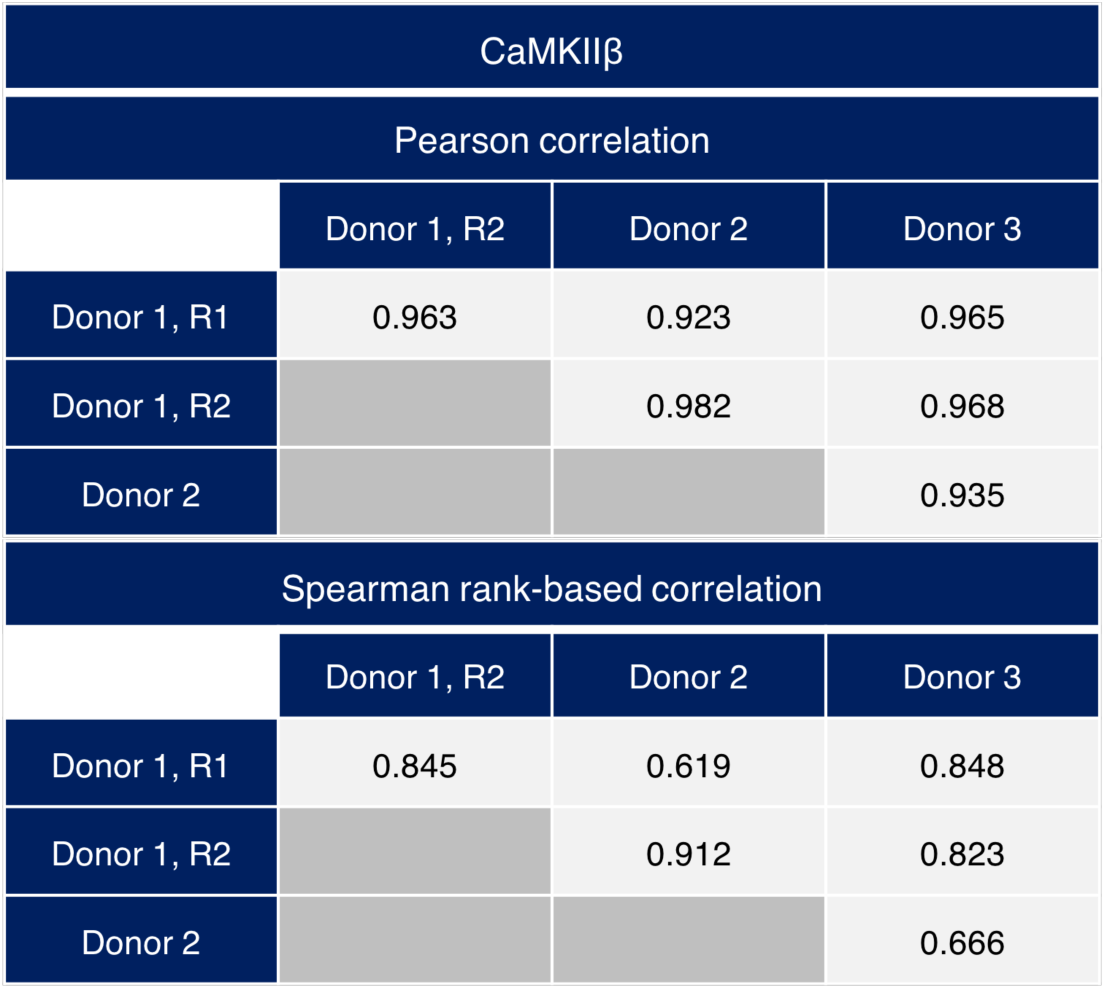
Pearson and Spearman correlation matrices for CaMKIIβ variants

**Table S18.** Pearson and Spearman correlation matrices for CaMKIIγ variants

| CaMKIIγ | | | |
| --- | --- | --- | --- |
| Pearson correlation |  |  |  |
|  | Donor 1, R2 | Donor 2 | Donor 3 |
| Donor 1, R1 | 0.913 | 0.771 | 0.498 |
| Donor 1, R2 |  | 0.939 | 0.673 |
| Donor 2 |  |  | 0.535 |
| Spearman rank-based correlation |  |  |  |
|  | Donor 1, R2 | Donor 2 | Donor 3 |
| Donor 1, R1 | 0.870 | 0.608 | 0.733 |
| Donor 1, R2 |  | 0.891 | 0.874 |
| Donor 2 |  |  | 0.767 |

**Table S19.** Pearson and Spearman correlation matrices for CaMKIIδ variants

| CaMKIIδ | | | |
| --- | --- | --- | --- |
| Pearson correlation |  |  |  |
|  | Donor 1, R2 | Donor 2 | Donor 3 |
| Donor 1, R1 | 0.974 | 0.927 | 0.823 |
| Donor 1, R2 |  | 0.932 | 0.924 |
| Donor 2 |  |  | 0.841 |
| Spearman rank-based correlation |  |  |  |
|  | Donor 1, R2 | Donor 2 | Donor 3 |
| Donor 1, R1 | 0.855 | 0.965 | 0.846 |
| Donor 1, R2 |  | 0.931 | 0.912 |
| Donor 2 |  |  | 0.949 |

**File S1.** Exon sequences for all CaMKII genes

Italicized nucleotides indicate optionally excluded portions of exons

CaMKIIα exons sequences

>Exon 1 (coding part only) ATGGCCACCATCACCTGCACCCGCTTCACGGAAGAGTACCAGCTCTTCGAGGAATTGGGCAA

>Exon 2 GGGAGCCTTCTCGGTGGTGCGAAGGTGTGTGAAGGTGCTGGCTGGCCAGGAGTATGCTGCCAAGATCATCAACACAAAGAAGCTGTCAGCCAGAG

>Exon 3 ACCATCAGAAGCTGGAGCGTGAAGCCCGCATCTGCCGCCTGCTGAAGCACCCCAACATCG

>Exon 4 TCCGACTACATGACAGCATCTCAGAGGAGGGACACCACTACCTGATCTTCGACCT

>Exon 5 GGTCACTGGTGGGGAACTGTTTGAAGATATCGTGGCCCGGGAGTATTACAGTGAGGCGGATGCCAG

>Exon 6 TCACTGTATCCAGCAGATCCTGGAGGCTGTGCTGCACTGCCACCAGATGGGGGTGGTGCACCGGGACCTGAAG

>Exon 7 CCTGAGAATCTGTTGCTGGCCTCCAAGCTCAAGGGTGCCGCAGTGAAGCTGGCAGACTTTGGCCTGGCCATAGAGGTGGAGGGGGAGCAGCAGGCATGGTTTG

>Exon 8 GGTTTGCAGGGACTCCTGGATATCTCTCCCCAGAAGTGCTGCGGAAGGACCCGTACGGGAAGCCTGTGGACCTGTGGGCTTGTG

>Exon 9 GGGTCATCCTGTACATCCTGCTGGTTGGGTACCCCCCGTTCTGGGATGAGGACCAGCACCGCCTGTACCAGCAGATCAAAGCCGGCGCCTATGAT

>Exon 10 TTCCCATCGCCGGAATGGGACACTGTCACCCCGGAAGCCAAGGATCTGATCAATAAGATGCTGACCATTAACCCATCCAAACGCATCACAGCTGCCGAAGCCCTTAAGCACCCCTGGATCTCG

>Exon 11 CACCGCTCCACCGTGGCATCCTGCATGCACAGACAGGAGACCGTGGACTGCCTGAAGAAGTTCAATGCCAGGAGGAAACTGAAG

>Exon 12 GGAGCCATTCTCACCACGATGCTGGCCACCAGGAACTTCTCCG

>Exon 14 GAGGGAAGAGTGGGGGAAACAAGAAGAGCGATGGTGTGAA

>Exon 15 GAAAAGAAAGTCCAGTTCCAGCGTTCAGTTAAT

>Exon 18 GGAATCCTCAGAGAGCACCAACACCACCATCGAGGATGAAGACACCAAAG

>Exon 22 TGCGGAAACAGGAAATTATAAAAGTGACAGAGCAGCTGATTGAAGCCATAAGCAATGGAGATTTTGAGTCCTACAC

>Exon 23 GAAGATGTGCGACCCTGGCATGACAGCCTTCGAACCTGAGGCCCTGGGGAACCTGGTTGAGGGCCTGGACTTCCATCGATTCTATTTTGAAAACC

>Exon 24 TGTGGTCCCGGAACAGCAAGCCCGTGCACACCACCATCCTGAATCCCCACATCCACCTGATGGGCGACGAGTCAGCCTGCATCGCCTACATCCGCATCACGCAGTACCTGGACGCTGGCGGCATCCCACGCACCGCCCAGTCGGAGGAGACCCGTGTCTGGCACCGCCGGGATGGCAAATGGCAGATCGTCCACTTCCACAGATCTGGGGCGCCCTCCGTCCTGCCCCA

>Exon 25 (coding part only) CTGA

CaMKIIβ exon sequences

>Exon 1 (coding part only) ATGGCCACCACGGTGACCTGCACCCGCTTCACCGACGAGTACCAGCTCTACGAGGATATTGGCAA

>Exon 2 GGGGGCTTTCTCTGTGGTCCGACGCTGTGTCAAGCTCTGCACCGGCCATGAGTATGCAGCCAAGATCATCAACACCAAGAAGCTGTCAGCCAGAG

>Exon 3 ATCACCAGAAGCTGGAGAGAGAGGCTCGGATCTGCCGCCTTCTGAAGCATTCCAACATCG

>Exon 4 TGCGTCTCCACGACAGCATCTCCGAGGAGGGCTTCCACTACCTGGTCTTCGATCT

>Exon 5 GGTCACTGGTGGGGAGCTCTTTGAAGACATTGTGGCGAGAGAGTACTACAGCGAGGCTGATGCCAG

>Exon 6 TCACTGTATCCAGCAGATCCTGGAGGCCGTTCTCCATTGTCACCAAATGGGGGTCGTCCACAGAGACCTCAAG

>Exon 7 CCGGAGAACCTGCTTCTGGCCAGCAAGTGCAAAGGGGCTGCAGTGAAGCTGGCAGACTTCGGCCTAGCTATCGAGGTGCAGGGGGACCAGCAGGCATGGTTTG

>Exon 8 GTTTCGCTGGCACACCAGGCTACCTGTCCCCTGAGGTCCTTCGCAAAGAGGCGTATGGCAAGCCTGTGGACATCTGGGCATGTG

>Exon 9 GGGTGATCCTGTACATCCTGCTCGTGGGCTACCCACCCTTCTGGGACGAGGACCAGCACAAGCTGTACCAGCAGATCAAGGCTGGTGCCTATGAC

>Exon 10 TTCCCGTCCCCTGAGTGGGACACCGTCACTCCTGAAGCCAAAAACCTCATCAACCAGATGCTGACCATCAACCCTGCCAAGCGCATCACAGCCCATGAGGCCCTGAAGCACCCGTGGGTCTGC

>Exon 11 CAACGCTCCACGGTAGCATCCATGATGCACAGACAGGAGACTGTGGAGTGTCTGAAAAAGTTCAATGCCAGGAGAAAGCTCAAG

>Exon 12 GGAGCCATCCTCACCACCATGCTGGCCACACGGAATTTCTCAG

>Exon 13 TGGGCAGACAGACCACCGCTCCGGCCACAATGTCCACCGCGGCCTCCGGCACCACCATGGGGCTGGTGGAACAAG

>Exon 14

*CAG*CCAAGAGTTTACTCAACAAGAAAGCAGATGGAGTCAAG

>Exon 16 CCCCAGACGAATAGCACCAAAAACAGTGCAGCCGCCACCAGCCCCAAAGGGACGCTTCCTCCTGCCGCCCTG

>Exon 17 GAGCCTCAAACCACCGTCATCCATAACCCAGTGGACGGGATTAAG

>Exon 18 GAGTCTTCTGACAGTGCCAATACCACCATAGAGGATGAAGACGCTAAAG

>Exon 19 CCCCCAGGGTCCCCGACATCCTGAGCTCAGTGAGGAGGGGCTCGGGAGCCCCAGAAGCCGAGGGGCCCCTGCCCTGCCCATCTCCGGCTCCCTTTAGCCCCCTGCCAGCCCCAT

>Exon 20 CCCCCAGGATCTCTGACATCCTGAACTCTGTGAGAAGGGGTTCAGGAACCCCAGAAGCCGAGGGCCCCCTCTCAGCGGGGCCCCCGCCCTGCCTGTCTCCGGCTCTCCTAGGCCCCCTGTCCTCCCCGT

>Exon 21 CCCCCAGGATCTCTGACATCCTGAACTCTGTGAGGAGGGGCTCAGGGACCCCAGAAGCCGAGGGCCCCTCGCCAGTGGGGCCCCCGCCCTGCCCATCTCCGACTATCCCTGGCCCCCTGCCCACCCCAT

>Exon 22 CCCGGAAGCAGGAGATCATTAAGACCACGGAGCAGCTCATCGAGGCCGTCAACAACGGTGACTTTGAGGCCTACGC

>Exon 23 *GAAAATCTGTGACCCAGGGCTGACCTCGTTTGAGCCTGAAGCACTGGGCAACCTGGTTGAAGGGATGG ACTTCCACAG*ATTCTACTTCGAGAACC

>Exon 24 TGCTGGCCAAGAACAGCAAGCCGATCCACACGACCATCCTGAACCCACACGTGCACGTCATTGGAGAGGATGCCGCCTGCATCGCTTACATCCGGCTCACGCAGTACATTGACGGGCAGGGCCGGCCCCGCACCAGCCAGTCTGAGGAGACCCGCGTGTGGCACCGCCGCGACGGCAAGTGGCAGAACGTGCACTTCCACTGCTCGGGCGCGCCTGTGGCCCCGCTGCAGTGA

CaMKIIγ exon sequences

>Exon 1 (coding part only) ATGGCCACCACCGCCACCTGCACCCGTTTCACCGACGACTACCAGCTCTTCGAGGAGCTTGGCAA

>Exon 2 GGGTGCTTTCTCTGTGGTCCGCAGGTGTGTGAAGAAAACCTCCACGCAGGAGTACGCAGCAAAAATCATCAATACCAAGAAGTTGTCTGCCCGGG

>Exon 3 ATCACCAGAAACTAGAACGTGAGGCTCGGATATGTCGACTTCTGAAACATCCAAACATCG

>Exon 4 TGCGCCTCCATGACAGTATTTCTGAAGAAGGGTTTCACTACCTCGTGTTTGACCT

>Exon 5 TGTTACCGGCGGGGAGCTGTTTGAAGACATTGTGGCCAGAGAGTACTACAGTGAAGCAGATGCCAG

>Exon 6 CCACTGTATACATCAGATTCTGGAGAGTGTTAACCACATCCACCAGCATGACATCGTCCACAGGGACCTGAAG

>Exon 7 CCTGAGAACCTGCTGCTGGCGAGTAAATGCAAGGGTGCCGCCGTCAAGCTGGCTGATTTTGGCCTAGCCATCGAAGTACAGGGAGAGCAGCAGGCTTGGTTTG

>Exon 8 GTTTTGCTGGCACCCCAGGTTACTTGTCCCCTGAGGTCTTGAGGAAAGATCCCTATGGAAAACCTGTGGATATCTGGGCCTGCG

>Exon 9 GGGTCATCCTGTATATCCTCCTGGTGGGCTATCCTCCCTTCTGGGATGAGGATCAGCACAAGCTGTATCAGCAGATCAAGGCTGGAGCCTATGAT

>Exon 10 TTCCCATCACCAGAATGGGACACGGTAACTCCTGAAGCCAAGAACTTGATCAACCAGATGCTGACCATAAACCCAGCAAAGCGCATCACGGCTGACCAGGCTCTCAAGCACCCGTGGGTCTGT

>Exon 11 CAACGATCCACGGTGGCATCCATGATGCATCGTCAGGAGACTGTGGAGTGTTTGCGCAAGTTCAATGCCCGGAGAAAACTGAAG

>Exon 12 GGTGCCATCCTCACGACCATGCTTGTCTCCAGGAACTTCTCAG

>Exon 13 TTGGCAGGCAGAGCTCCGCCCCCGCCTCGCCTGCCGCGAGCGCCGCCGGCCTGGCCGGGCAAG

>Exon 14 CTGCCAAAAGCCTATTGAACAAGAAGTCGGATGGCGGTGTCAAG

>Exon 15 AAAAGGAAGTCGAGTTCCAGCGTGCACCTAATG

>Exon 16 CCACAGAGCAACAACAAAAACAGTCTCGTAAGCCCAGCCCAAGAGCCCGCGCCCTTGCAGACGGC CATG

>Exon 17 GAGCCACAAACCACTGTGGTACACAACGCTACAGATGGGATCAAG

>Exon 18 GGCTCCACAGAGAGCTGCAACACCACCACAGAAGATGAGGACCTCAAAG*GTAG*

>Exon 19 *CTGCCCCGCTCCGCACTGGGAATGGCAGCTC*GGTGCCTGAAGGACGGAGCTCCCGGGACAGAACAG CCCCCTCTGCAGGCATGCAGCCCCAGCCTTCTCTCTGCTCCTCAGCCA

>Exon 22 TGCGAAAACAGGAGATCATTAAGATTACAGAACAGCTGATTGAAGCCATCAACAATGGGGACTTTGAGGCCTACAC

>Exon 23 GAAGATTTGTGATCCAGGCCTCACTTCCTTTGAGCCTGAGGCCCTTGGTAACCTCGTGGAGGGGATGGATTTCCATAAGTTTTACTTTGAGAATC

>Exon 24 (coding part only) TCCTGTCCAAGAACAGCAAGCCTATCCATACCACCATCCTAAACCCACACGTCCACGTGATTGGGGAGGACGCAGCGTGCATCGCCTACATCCGCCTCACCCAGTACATCGACGGGCAGGGTCGGCCTCGCACCAGCCAGTCAGAAGAGACCCGGGTCTGGCACCGTCGGGATGGCAAGTGGCTCAATGTCCACTATCACTGCTCAGGGGCCCCTGCCGCACCGCTGCAGTGA

CaMKIIδ exon sequences

>Exon 1 (coding part only) ATGGCTTCGACCACAACCTGCACCAGGTTCACGGACGAGTATCAGCTTTTCGAGGAGCTTGGAAA

>Exon 2 GGGGGCATTCTCAGTGGTGAGAAGATGTATGAAAATTCCTACTGGACAAGAATATGCTGCCAAAATTATCAACACCAAAAAGCTTTCTGCTAGGG

>Exon 3 ATCATCAGAAACTAGAAAGAGAAGCTAGAATCTGCCGTCTTTTGAAGCACCCTAATATTG

>Exon 4 TGCGACTTCATGATAGCATATCAGAAGAGGGCTTTCACTACTTGGTGTTTGATTT

>Exon 5 AGTTACTGGAGGTGAACTGTTTGAAGACATAGTGGCAAGAGAATACTACAGTGAAGCTGATGCCAG

>Exon 6v1 TCATTGTATACAGCAGATTCTAGAAAGTGTTAATCATTGTCACCTAAATGGCATAGTTCACAGGGACCTGAAG

>Exon 6v2 TCATTGCATTCAGCAGATCCTGGAGGCTGTGCTACACTGCCATCAGATGGGCGTGGTCCATCGGGACCTGAAG

>Exon 7 CCTGAGAATTTGCTTTTAGCTAGCAAATCCAAGGGAGCAGCTGTGAAATTGGCAGACTTTGGCTTAGCCATAGAAGTTCAAGGGGACCAGCAGGCGTGGTTTG

>Exon 8 GTTTTGCTGGCACACCTGGATATCTTTCTCCAGAAGTTTTACGTAAAGATCCTTATGGAAAGCCAGTGGATATGTGGGCATGTG

>Exon 9 GTGTCATTCTCTATATTCTACTTGTGGGGTATCCACCCTTCTGGGATGAAGACCAACACAGACTCTATCAGCAGATCAAGGCTGGAGCTTATGAT

>Exon 10 TTTCCATCACCAGAATGGGACACGGTGACTCCTGAAGCCAAAGACCTCATCAATAAAATGCTTACTATCAACCCTGCCAAACGCATCACAGCCTCAGAGGCACTGAAGCACCCATGGATCTGT

>Exon 11 CAACGTTCTACTGTTGCTTCCATGATGCACAGACAGGAGACTGTAGACTGCTTGAAGAAATTTAATGCTAGAAGAAAACTAAAG

>Exon 12 GGTGCCATCTTGACAACTATGCTGGCTACAAGGAATTTCTCAG

>Exon 14

*CAG*CCAAGAGTTTGTTGAAGAAACCAGATGGAGTAAAG

>Exon 15 AAAAGGAAGTCCAGTTCGAGTGTTCAGATGATG

>Exon 16 ATAAACAACAAAGCCAACGTGGTAACCAGCCCCAAAGAAAATATTCCTACCCCAGCGCTG

>Exon 17 GAGCCCCAAACTACTGTAATCCACAACCCTGATGGAAACAAG

>Exon 18 GAGTCAACTGAGAGTTCAAATACAACAATTGAGGATGAAGATGTGAAAG

>Exon 22 CACGAAAGCAAGAGATTATCAAAGTCACTGAACAACTGATCGAAGCTATCAACAATGGGGACTT TGAAGCCTACAC

>Exon 23 AAAAATCTGTGACCCAGGCCTTACTGCTTTTGAACCTGAAGCTTTGGGTAATTTAGTGGAAGGG ATGGATTTTCACCGATTCTACTTTGAAAATG

>Exon 24 CTTTGTCCAAAAGCAATAAACCAATCCACACTATTATTCTAAACCCTCATGTACATCTGGTAGGG GATGATGCCGCCTGCATAGCATATATTAGGCTCACACAGTACATGGATGGCAGTGGAATGCCAA AGACAATGCAGTCAGAAGAGACTCGTGTGTGGCACCGCCGGGATGGAAAGTGGCAGAATGTTCA TTTTCATCGCTCGGGGTCACCAACAGTACCCATCAA

>Exon 25 (coding part only) GCCACCCTGTATTCCAAATGGGAAAGAAAACTTCTCAGGAGGCACCTCTTTGTGGCAAAACATCT AA

>Exon 26 (coding part only) GTAA

**File S2.** Primers for initial amplification of variably spliced transcript regions CaMKIIα

F: CCTGGATCTCGCACCGCTCCACCGT

R: GGCTGTCATGCCAGGGTCGCACATCTTCG

CaMKIIβ

F: CCTGAGGTCCTTCGCAAAGAGGCGTATGGC R: GCAGGCGGCATCCTCTCCAATGACGTGC

CaMKIIγ

F: GCGCATCACGGCTGACCAGGCTCTCA

R: CCATCCCCTCCACGAGGTTACCAAGGGCC

CaMKIIδ

F: GCTTGGAAAGGGGGCATTCTCAGTGGTGAGAAGATG R: TGCAGGCGGCATCATCCCCTACCAGATGTACA

Gene-specific primers for Illumina sequencing library construction:

CaMKIIα

F: CATCCTCACCACCATGCTG R: CCAGCAGGTTCTCGAAGTAG

CaMKIIβ

F: TGGATCTCGCACCGCT

R: GTCGCACATCTTCGTGTAGG

CaMKIIγ

F: CCTCACGACCATGCTTGTCT R: CCCCTCCACGAGGTTACC

CaMKIIδ

F: CAGTGAAGCTGATGCCAGTC

R: TGTTCAGTGACTTTGATAATCTCTTGC

Library construction primers contain the following generic 3’ overhangs F: ACACTCTTTCCCTACACGACGCTCTTCCGATCT

R: CTGGAGTTCAGACGTGTGCTCTTCCGATC

